# *C. elegans* huntingtin, *htt-1,* promotes robust autophagy induction and survival under stress conditions

**DOI:** 10.1101/2024.12.01.625591

**Authors:** Christine Hyein Chung, Hanee Lee, Kyumin Park, Young Seo Park, Roy Jung, Ihnsik Seong, Junho Lee

## Abstract

Huntingtin (HTT) is the gene responsible for Huntington’s disease (HD), a neurodegenerative disorder caused by a CAG trinucleotide repeat expansion mutation. While HD pathogenesis has traditionally been attributed to the toxic gain-of-function effects of mutant huntingtin (mHTT), increasing evidence underscores the critical role of wild-type HTT loss-of-function. Understanding the physiological roles of HTT is essential for elucidating HD mechanisms and developing effective therapeutic strategies. The *C. elegans htt-1* gene, an ortholog of human HTT, remains largely uncharacterized. Here, we demonstrate that *htt-1* promotes survival under stress conditions that requires autophagy as a defense mechanism. Specifically, we identify intestinal *htt-1* as a key regulator of *C. elegans* survival during *Pseudomonas aeruginosa* PA14 infection. Our findings reveal that *htt-1* functions downstream of the MPK-1/ERK pathway to induce systemic autophagy and enhance host defense during immune challenges. Moreover, expression of wild-type human HTT in *htt-1* mutant worms rescues the survival defect during PA14 infection. Expression of mutant human HTT, on the other hand, exacerbates survival deficits, underscoring the conserved function of HTT across species. Additionally, we found that *htt-1* affects survival and autophagy under heat shock stress conditions. These results establish *htt-1* as a critical regulator of survival and autophagy in response to both pathogenic bacterial infection and thermal stress.

**Author summary:** Huntingtin plays essential roles in selective autophagy across various organisms, including *Drosophila,* mice, and humans. Notably, the *C. elegans* huntingtin ortholog, *htt-1,* remains largely uncharacterized. Here, we describe a novel pro-survival role *htt-1* in stress resistance. We show that *htt-1* mutants exhibit significantly reduced survival during *Pseudomonas aeruginosa* infection. Expression of wild-type human HTT rescues the reduced survival of *htt-1* mutants, whereas mutant HTT further exacerbates the phenotype. These findings establish *C. elegans htt-1* as a valuable model for studying huntingtin biology and its roles in stress resistance. Downregulation of ERK in *htt-1* mutants had no additional effect on survival, suggesting that *htt-1* functions downstream of ERK. Interestingly, *htt-1* mutants display reduced autophagy, indicating that *htt-1* functions in the ERK-autophagy signaling cascade during infection. Mechanistically, we propose that *htt-1* regulates autophagy at the protein level. HTT-1 remains cytoplasmic, and autophagy-related gene transcripts remain stable, without downregulation, in *htt-1* mutants following infection. Additionally, *htt-1* mutants exhibit reduced survival and impaired autophagy under heat stress. These results suggest that *htt-1* may have a broader role in stress resilience extending beyond immune defense.

## Introduction

Huntington disease (HD) is an autosomal dominant neurodegenerative disorder that affects approximately 5 to 10 per 10,000 individuals in the Caucasian population. Clinically, HD is characterized by motor disturbances, cognitive decline, and psychiatric symptoms. A hallmark of HD pathology is the aggregation of neuronal proteins, which causes morphological and functional defects, ultimately leading to neuronal death, particularly in the striatum [1]. The huntingtin (HTT) gene, which is linked to HD, encodes a large, ubiquitously expressed 348 kDa protein. The aggregates and cellular dysfunction are driven by the CAG glutamine repeat expansion mutation near the N-terminus of the HTT gene. These hallmark features are observed not only in HD patients but also in *C. elegans* and other animal HD models. For example, expressing human mutant HTT (mHTT) exon 1 fragments in *C. elegans* led to apoptotic cell death in ASH sensory neurons and morphological and functional abnormalities in PLM sensory neurons [2,3]. Additionally, human mHTT expression in body wall muscle cells caused motility defects, mitochondrial fragmentation, and disruption of the mitochondrial network in *C. elegans* [4-6]. The toxicity of mHTT and its contribution to pathology are widely studied. Furthermore, emerging evidence suggests that the loss of wild-type HTT function may also contribute to HD pathogenesis.

Although wild-type HTT is known to be involved in several cellular processes, including vesicular transport, gene transcription, and autophagy in specific experimental conditions [1], only limited information is available for normal functions of HTT at the level of genes and organism. We decided to utilize the molecular genetic tools of the model organism *C. elegans* because it contains an ortholog of human huntingtin (HTT), the *htt-1* gene, which has remained largely uncharacterized. The *htt-1* gene is located on the X chromosome and produces six transcript isoforms, with the primary isoform, HTT-1a, consisting of 2022 amino acids. Although *htt-1* lacks the polyglutamine (polyQ) tract present in human HTT, structural predictions suggest a high degree of structural homology between human Htt and two ordered regions of HTT-1, spanning amino acid sequences 1-1141 and 1241-2022. These regions show high TM scores (>0.8) and low RMSD values (≤2.0 Å) [7]. This structural conservation provides a valuable model for investigating the physiological functions of HTT in model organisms. Understanding the functions of HTT is essential for developing effective therapies for HD. Although treatments for HD primarily address behavioral symptoms, recent efforts have focused on developing disease-modifying therapies. Reducing mHTT expression holds promise, but the threshold at which normal or mutant HTT levels can be reduced without causing cellular dysfunction remains unclear [8].

Despite over 30 years since the discovery of the HD-causing gene, the importance of HTT’s normal function in cellular processes remains unclear. This study aims to establish *C. elegans* huntingtin ortholog *htt-1* as a viable model for studying huntingtin function. We demonstrate that *C. elegans htt-1* plays a key role in regulating autophagy via ERK activity during *Pseudomonas aeruginosa* PA14 infection. Our findings highlight the critical role of *htt-1* in promoting survival and enhancing autophagy in response to both pathogenic bacterial infection and thermal stress. Leveraging the conserved biology of HTT across species, this research aims to broaden our understanding of HTT functions and mechanisms.

## Results

### Increased mortality of *htt-1* mutants caused by *Pseudomonas aeruginosa* PA14 infection

The *C. elegans htt-1* gene exhibits broad expression across various tissues in both hermaphrodites and males throughout all developmental stages, including embryonic, larval, adult and dauer stages (Fig 1A). To investigate the function of *htt-1*, we employed reverse genetics to analyze two deletion mutants, *tm8121* and *tm1959*. The *tm8121* allele contains a 100 bp deletion within exon 17, resulting in a frameshift, while *tm1959* allele carries a 506 bp deletion spanning exons 2-4 and their adjacent intron sequences (Fig 1B). Despite the highest *htt-1* expression being detected during the embryonic stage, neither mutation affected development under standard laboratory conditions [9]. Both *tm8121* and *tm1959* mutants displayed normal morphology and growth. Additionally, brood size and lifespan measurements showed no significant differences between wild-type N2 and the *htt-1* mutant strains under standard culture conditions (Figs 1C and 1D). These findings indicate that *htt-1* is not essential for normal development, reproduction, or lifespan maintenance under optimal growth conditions.

**Fig 1.**
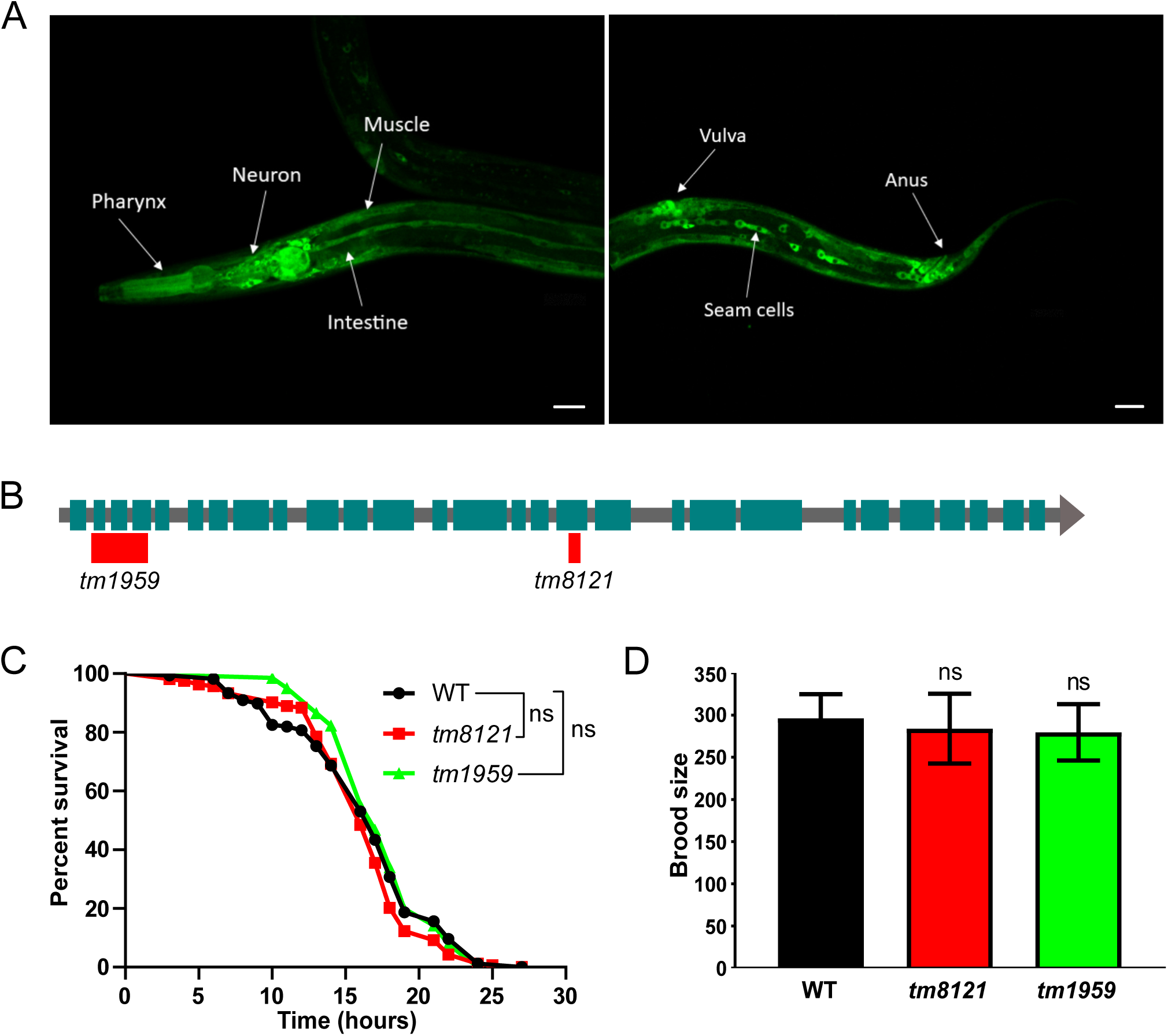
Mutation in the broadly expressed *htt-1* does not significantly affect lifespan or brood size. (A) Confocal image showing HTT-1::GFP expression in wild-type (WT) animals. (B) Schematic diagram of the *htt-1* gene structure. Teal boxes represent exons, black lines represent introns, and red boxes indicate the 506 bp deletion in *tm1959* and the 100 bp deletion in *tm8121.* (C) Lifespan analysis of WT and *htt-1* mutant animals (45-60 animals per replicate, *n* = 3). (D) Brood size analysis of WT and *htt-1* mutants (approximately 7 animals per replicate, *n* = 2). Error bars represent SD. Scale bars: 20 μm. Experiments were performed at 20_. Statistical significance was determined by the Log-rank test: ns, not significant (P > 0.05). Detailed statistical analyses for lifespan and brood size are provided in S1 Table.

*htt-1* mRNA levels increase after 24 hours of infection with pathogenic bacteria, such as *Pseudomonas aeruginosa* PA14 and *Photorhabdus luminescens,* compared to control conditions with nonpathogenic bacteria [10,11]. This observed upregulation suggests a potential role for *htt-1* in the immune response. To investigate this, we assessed the survival of synchronized L4 stage *C. elegans* transferred to pathogenic bacterium PA14. Both *htt-1* mutants, *tm8121* and *tm1959,* exhibited significantly reduced survival compared to wild-type N2 animals (Fig 2A). This survival defect was fully rescued by expressing wild-type *htt-1* under its own promoter (Fig 2B). To evaluate whether *htt-1* influences immune responses across different life stages, we assessed the survival of older adult worms exposed to PA14. Consistent with results observed in L4-stage animals, *htt-1(tm8121)* mutants showed increased susceptibility to PA14 at both day 1 and day 3 of adulthood (Figs 2C and 2D). These findings establish that *htt-1* is essential for *C. elegans* survival during PA14 infection, underscoring its critical role in host defense mechanisms across different life stages.

**Fig 2.**
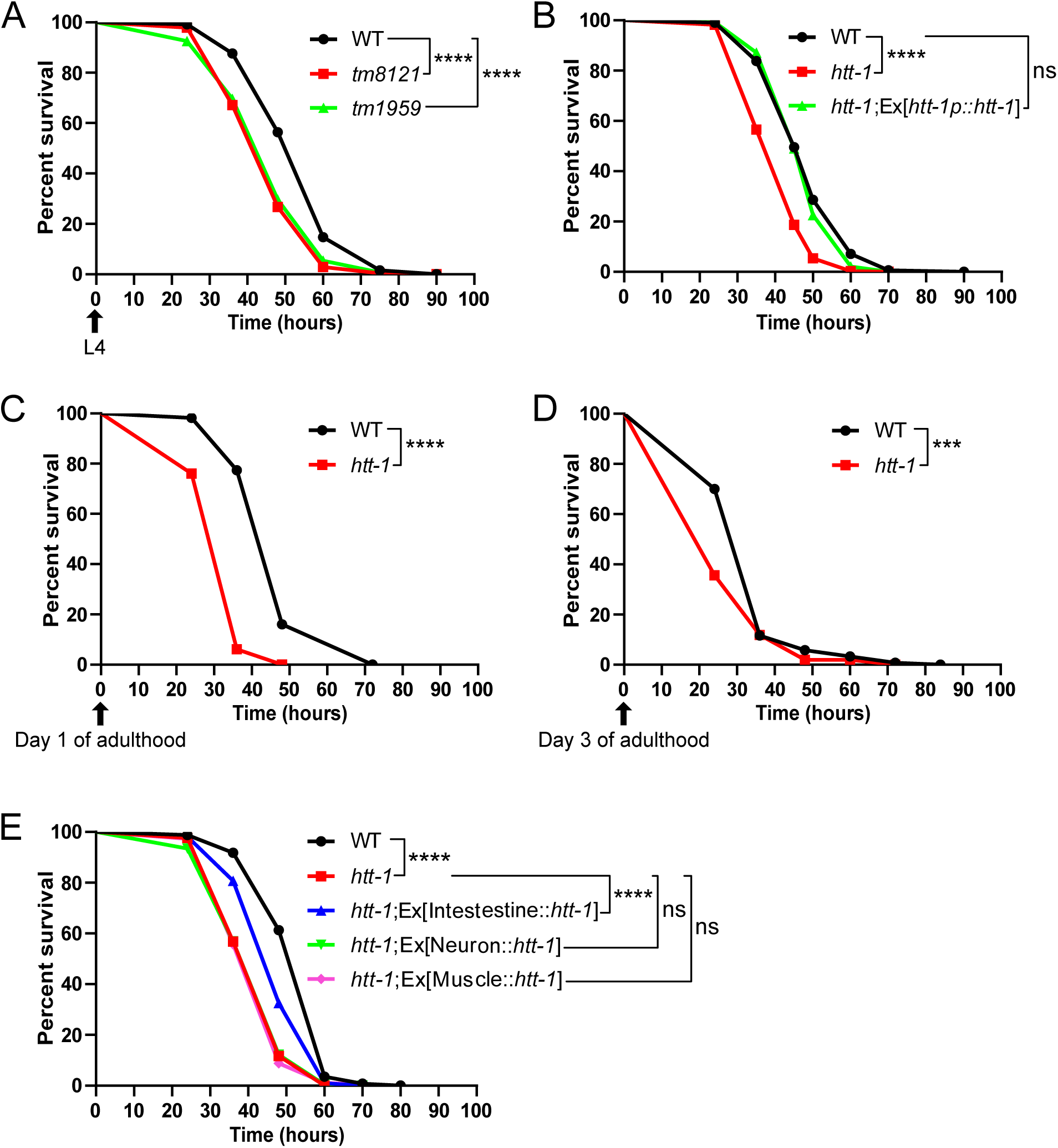
HTT-1 is essential for survival during PA14 infection across developmental stages. (A) Survival curves of L4-staged *C. elegans* wild-type N2 and *htt-1 (tm8121* and *tm1959)* mutant animals during PA14 infection (25-35 animals per replicate, *n* = 3). (B) Survival curves of *htt-1(tm8121)* mutant rescue experiments. Approximately 25-35 animals per replicate (*n* = 4), comparing wild-type (WT), *htt-1(tm8121)* mutants, and *htt-1(tm8121)*;Ex[*phtt-1::htt-1*] rescue animals. (C) Survival curves of day 1 adult N2 and *htt-1(tm8121)* mutant animals during PA14 infection (25-35 animals per replicate, *n* = 2). (D) Survival curves of day 3 adult N2 and *htt-1(tm8121)* mutant animals during PA14 infection (15-20 animals per replicate, *n* = 2). (E) Survival curves for tissue specific rescue of *htt-1(tm8121)* mutants during PA14 infection (25-35 animals per replicate, *n* = 3). Statistical significance was determined by the Log-rank test: ns, not significant (P > 0.05); *** *p* ≤ 0.001; **** *p* ≤ 0.0001. Detailed statistical analyses are provided in S1 Table.

Reduced survival of *htt-1* mutants during PA14 infection could result from differences in pathogen intake. To test this hypothesis, a lawn occupancy assay was performed to assess pathogen interaction in *htt-1(tm8121)* mutants and wild-type N2 animals. The test was conducted over 24 hours. No significant differences in lawn occupancy were observed during the first 8 hours, indicating that both strains displayed similar aversion behavior initially (S1 Fig). However, after 8 hours, *htt-1(tm8121)* mutants showed increased avoidance behavior compared to wild-type controls (S1 Fig). This shift in behavior is likely driven by intestinal lumen bloating, which activates neuroendocrine signaling to promote microbial aversion behavior [12]. The difference in lawn occupancy observed between 8 and 12 hours may reflect the emergence of physiological changes in *htt-1* mutants compared to wild-type N2, potentially influencing survival outcomes. However, the similar pathogen avoidance behavior during the first 8 hours suggests that the reduced survival is not due to differences in pathogen exposure between the two strains (S1 Fig).

Next, to investigate the molecular mechanism underlying *htt-1* function in response to infection, tissue specific rescue experiments were conducted based on the observed HTT-1 expression pattern (S2 Fig). Intestine-specific expression of wild-type *htt-1* under the *opt-2* promoter significantly rescued the mutant phenotype, improving survival during PA14 infection (Fig 2E). In contrast, expression of *htt-1* under the *myo-3* and *egl-3* promoters, which drive muscle- and neuron-specific expression respectively, failed to rescue the mutant phenotype and showed no significant survival difference compared to *htt-1(tm8121)* mutants (Fig 2E). These results highlight that HTT-1 action in the intestine is essential for its pro-survival role during PA14 infection. In *C. elegans*, intestinal epithelial cells form a primary defense line against ingested pathogens. Upon infection, key immune responses, including the ERK pathway and autophagy, are activated in the intestine [13]. However, while intestinal expression of *htt-1* was sufficient to enhance survival, it did not fully rescue the *htt-1(tm8121)* mutant phenotype. This suggests that HTT-1 activity in other tissues may also contribute to the organism’s overall survival during PA14 infection.

### Functional conservation of *htt-1* and human HTT in host defense during infection

Huntington’s disease (HD) is characterized by the expansion of polyQ repeats in the HTT gene, with a threshold of approximately 40 repeats for disease onset [14]. To investigate functional conservation, we generated transgenic *C. elegans* expressing either wild-type human HTT (with 26 polyQ repeats) or mutant human HTT (mHTT, with 76 polyQ repeats), both driven by the *C. elegans htt-1* promoter. Both strains showed no observable phenotype under standard laboratory conditions. However, when exposed to the pathogenic bacterium PA14, wild-type human HTT-expressing *htt-1(tm8121)* worms showed fully rescued survival, comparable to that of wild-type N2 animals (Fig 3). In contrast, worms expressing human mHTT displayed lower survival rates than both wild-type N2 and *htt-1(tm8121)* mutants (Fig 3). These results are consistent with previous *C. elegans* HD model studies, which show that mHTT expression causes severe toxicity in the worms, including protein aggregate formation and increased oxidative stress [2-6]. Rescue experiments with human HTT expression in *C. elegans* suggest that both *htt-1* and wild-type human HTT share a conserved pro-survival role in immune defense.

**Fig 3.**
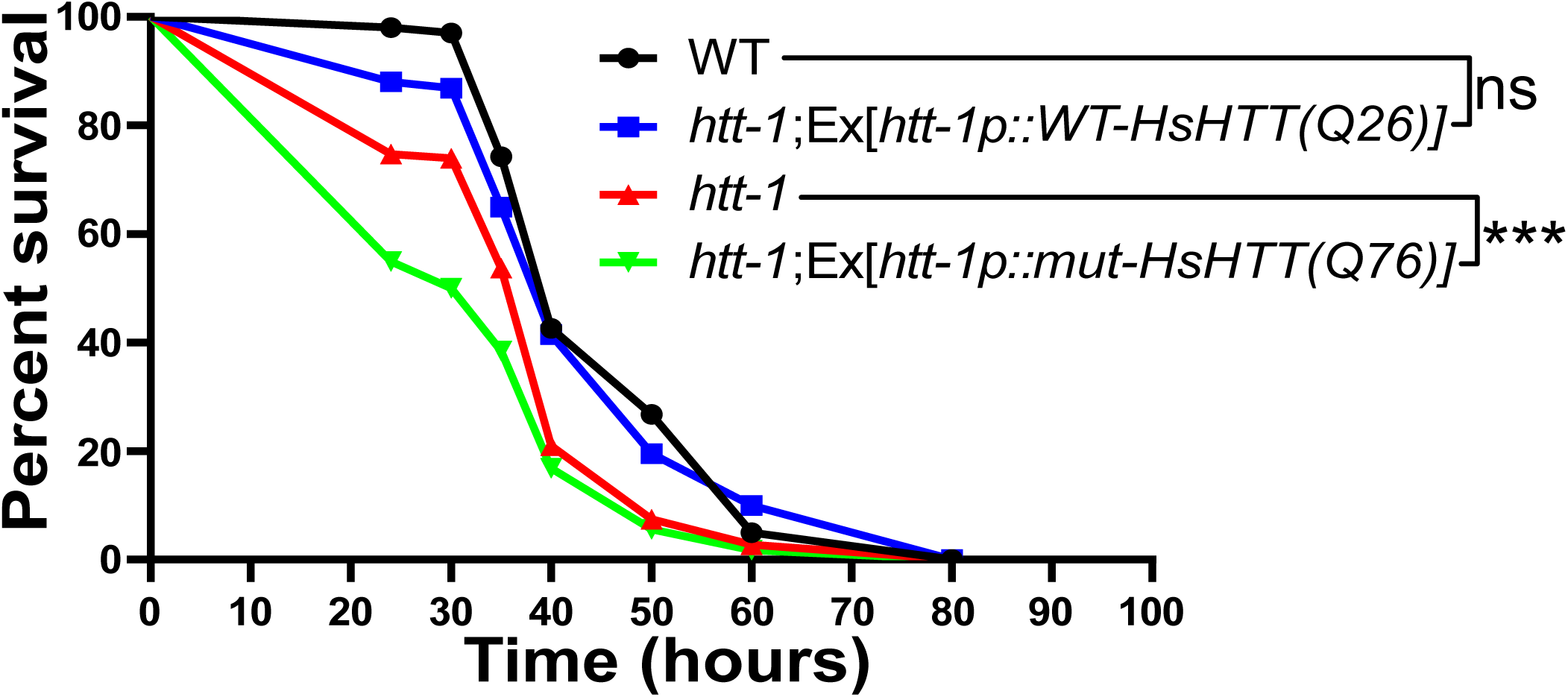
*htt-1* is functionally conserved with human huntingtin. Survival curves during PA14 infection showing rescue of *htt-1(tm8121)* by human HTT expression (25-35 animals per replicate, *n* = 3). Statistical significance was determined using the Log-rank test: ns, not significant (*p* > 0.05), *** *p* ≤ 0.001. Detailed statistical analyses of lifespan are provided in S1 Table.

### HTT-1 acts downstream of MPK-1/ERK pathway

To further investigate the molecular mechanisms underlying intestinal *htt-1* function in host defense during PA14 infection, we examined its involvement in potential pathways. A previous study demonstrated that autophagy-defective *C. elegans* exhibit decreased survival during PA14 infection due to increased necrosis activity. Notably, inhibiting necrosis through RNA interference (RNAi) targeting the cysteine peptidase *clp-1* enhances survival in autophagy-defective worms [13]. Based on these findings, we hypothesized that upregulated necrosis may similarly contribute to the reduced survival of *htt-1(tm8121)* mutants during PA14 infection. However, RNAi knockdown of *clp-1* in *htt-1(tm8121)* mutants did not significantly improve the survival of infected worms (S3A Fig). In addition to necrosis, apoptosis, another well-known form of programmed cell death, was also considered as a potential contributor to the increased mortality observed in *htt-1(tm8121)* mutant phenotype. We tested the effect of RNAi knockdown of *ced-3,* a cysteine protease essential for initiating apoptosis in *C. elegans,* on the survival of *htt-1(tm8121)* mutants [15]. *ced-3* knockdown did not affect survival rates in either wild-type N2 or *htt-1(tm8121)* mutant worms (S3B Fig). Together, these findings suggest that the reduced survival observed in *htt-1(tm8121)* mutant worms exposed to PA14 is not due to increased activity of either necrosis or apoptosis.

The underlying mechanisms may involve other cellular processes, such as immune or stress response pathways. Therefore, we aimed to identify immune response pathways associated with *htt-1*. In *C. elegans,* major innate immunity signaling pathways include the FOXO transcription factor DAF-16, protein kinase D DKF-2, the G protein Gqα EGL-30, the G protein-coupled receptor FSHR-1, the p38 mitogen-activated protein kinase (MAPK) PMK-1, and the extracellular signal-regulated kinase (ERK) MAPK MPK-1 [13]. To explore the relationship between *htt-1* and these immune signaling pathways, RNAi knockdowns were performed. Knockdowns of *daf-16, dkf-2, egl-30, fshr-1,* and *pmk-1* significantly reduced survival in both wild-type N2 and *htt-1(tm8121)* mutants during PA14 infection (S4A-E Fig). The additive effects observed in the survival of *htt-1(tm8121)* mutants suggest that theses pathways act independently of *htt-1*. This indicates that *htt-1* does not directly interact with or influence these pathways in the immune response to PA14 infection.

In contrast, while RNAi knockdowns of key components of the ERK pathway, including *lin-3* (EGF), *let-60* (RAS), *mek-2* (MEK), and *mpk-1* (ERK), decreased survival in wild-type N2, they had no significant effect on the survival of *htt-1(tm8121)* mutants (Fig 4A-D). This suggests that *htt-1* function in ERK-mediated immune defense signaling cascade. We further tested the effect of the *lin-3* by examining the *lin-3 (e1417)* mutation, which reduces ERK activation following exposure to PA14 infection [13]. The survival assay revealed reduced survival rates during PA14 infection in *lin-3(e1417)* mutants compared to wild-type N2 (Fig 4E). However, *htt-1(tm8121)* and *htt-1(tm8121); lin-3(e1417)* mutants showed no significant difference in survival, further supporting a functional relationship between *htt-1* and the ERK pathway (Fig 4E). Next, we used the MEK inhibitor U0126, which reduces ERK activity in PA14 infected worms [13]. Addition of 8 μM U0126 to the PA14-seeding slow killing assay nematode growth medium reduced survival in wild-type N2 worms but had no effect on the survival rate of *htt-1(tm8121)* worms during infection (Fig 4F). These ERK reduction-of-function experiments suggest that *htt-1* is involved in the ERK-mediated immune response to PA14 infection.

**Fig 4.**
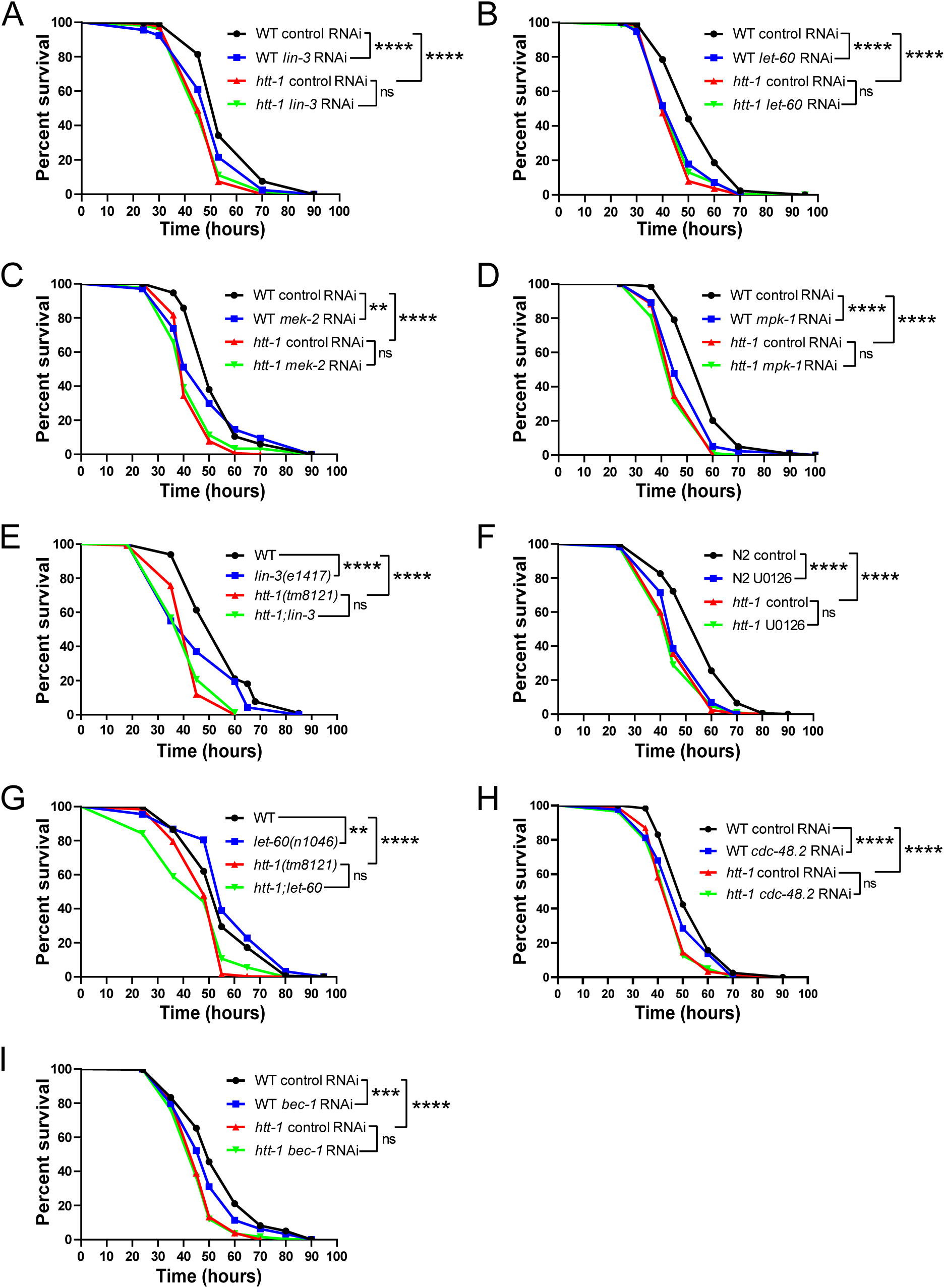
*htt-1* functions in the ERK-autophagy signaling pathways to promote survival during PA14 infection. (A) Percent survival of wild-type (WT) and *htt-1(tm8121)* animals with empty vector L4440 or *lin-3* RNAi during PA14 infection (25-35 animals per replicate, *n* = 3). (B) Percent survival of WT and *htt-1(tm8121)* animals with empty vector L4440 or *let-60* RNAi during PA14 infection (25-35 animals per replicate, *n* = 3). (C) Percent survival of WT and *htt-1(tm8121)* animals with empty vector L4440 or *mek-2* RNAi during PA14 infection (25-35 animals per replicate, *n* = 3). (D) Percent survival of WT and *htt-1(tm8121)* animals with empty vector L4440 or *mpk-1* RNAi during PA14 infection (25-35 animals per replicate, *n* = 3). (E) Percent survival of WT and *lin-3(e1417)* mutant animals (25-35 animals per replicate, *n* = 3). (F) Percent survival of WT and *htt-1(tm8121)* with or without U0126 (8 μM) treatment during PA14 infection (25-35 animals per replicate, *n* = 3). (G) Percent survival of WT and *let-60(n1046)* mutant animals (25-35 animals per replicate, *n* = 3). (H) Percent survival of WT and *htt-1(tm8121)* animals with empty vector L4440 or *cdc-48.2* RNAi during PA14 infection. (25-35 animals per replicate, *n* = 3). (I) Percent survival of WT and *htt-1(tm8121)* animals with empty vector L4440 or *bec-1* RNAi during PA14 infection. (25-35 animals per replicate, *n* = 3). ns (not significant) *p* > 0.05, ** *p* ≤ 0.01, *** *p* ≤ 0.001, **** *p* ≤ 0.0001 as determined by Log rank test. Detailed statistical analyses are provided in S1 Table.

To further explore the relationship between *htt-1* and the ERK pathway, we performed epistasis analysis. The *let-60/*RAS*(n1046)* gain-of-function mutation results in a constitutively active Ras protein, leading to upregulation of the ERK pathway in *C. elegans* [16]. The *let-60(n1046)* mutation increased survival in wild-type N2 worms during PA14 infection (Fig 4G). However, no significant survival difference was observed between *htt-1(tm8121)* and *htt-1(tm8121);let-60(n1046)* mutants (Fig 4G). These results indicate that functional *htt-1* is essential for the increased survival phenotype of *let-60(n1046)* mutants and that *htt-1* acts downstream of *let-60* in response to PA14 infection. Together, these results demonstrate that *let-60* and subsequent *htt-1* activity promote *C. elegans* survival during PA14 infection.

### *C. elegans cdc-48.2, htt-1,* and *bec-1* act downstream of ERK signaling in response to PA14 infection

In *C. elegans,* PA14 infection activates the intestinal ERK pathway, which leads to the phosphorylation of CDC-48.2 [13]. *cdc-48.2,* the *C. elegans* homolog of the AAA+ ATPase VCP/Cdc48/p97, is expressed in the intestine and plays a critical role in regulating ubiquitinated proteins and various cellular processes, including autophagy [17-19]. VCP is a highly conserved protein with homologs across species, including CDC-48.2 in *C. elegans*, VCP in humans, VCP in *Mus musculus,* TER97 in *Drosophila melanogaster,* and Cdc48 in *Saccharomyces cerevisiae* [20]. To explore a potential functional relationship between *htt-1* and *cdc-48.2*, RNAi knockdown of *cdc-48.2* was performed. *cdc-48.2* knockdown increased sensitivity to PA14 infection in wild-type N2 worms, but it did not significantly affect survival in *htt-1(tm8121)* mutant worms (Fig 4H).

In response to PA14 infection, activated CDC-48.2 triggers autophagy in *C. elegans* [13]. In human cells, VCP/p97 interacts with Beclin-1 to initiate autophagy [21]. To further investigate the role of *htt-1* in the ERK-mediated stress response during PA14 infection, RNAi knockdown of the autophagy gene *bec-1* was performed. *bec-1*, the *C. elegans* ortholog of yeast and mammalian autophagy proteins Atg6/Vps30 and Beclin1, is essential for development. Inactivation of *bec-1* activates the *ced-3* caspase-mediated cell death pathway, and strong loss-of-function or null alleles of *bec-1* are highly lethal [22]. Incomplete inactivation of *bec-1* through RNAi knockdown reduces, but does not fully eliminate, *bec-1* function, as demonstrated by reduced GFP fluorescence in worms expressing GFP-tagged BEC-1 [22]. Knockdown of *bec-1* decreased survival in wild-type N2 worms exposed to PA14, but did not further reduce survival in *htt-1(tm8121)* mutants (Fig 4I). These results suggest that *htt-1, cdc-48.2,* and *bec-1* may function within the same signaling cascade to promote survival during PA14 infection in *C. elegans*.

### *htt-1* induces robust autophagy in response to infection and heat stress

Results from the reduction-of-function experiments suggest that *htt-1* may play a role within the ERK-autophagy signaling pathways during PA14 infection in *C. elegans.* Autophagy is a critical pro-survival mechanism that enhances resistance to various stressors, including pathogen infection, heat shock, osmotic stress, oxidative stress, and starvation [23,24]. In *C. elegans,* autophagy activity is assessed *in vivo* by quantifying GFP::LGG-1 puncta, which label autophagosomes. LGG-1, the ortholog of Atg8/LC3, is a ubiquitin-like, microtubule-associated protein that facilitates autophagic vesicle growth [25]. Autophagic activity was analyzed in wild-type N2 and *htt-1(tm8121)* mutants within the first four intestinal cells (int1) and in seam cells, a population of epidermal stem cells. Both strains were exposed to either PA14 pathogen or nonpathogenic OP50 bacteria for 12 hours at 25°C. Upon PA14 infection, GFP::LGG-1 foci increased in wild-type N2 within seam cells and int1 cells (Fig 5A and 5C).

**Fig 5.**
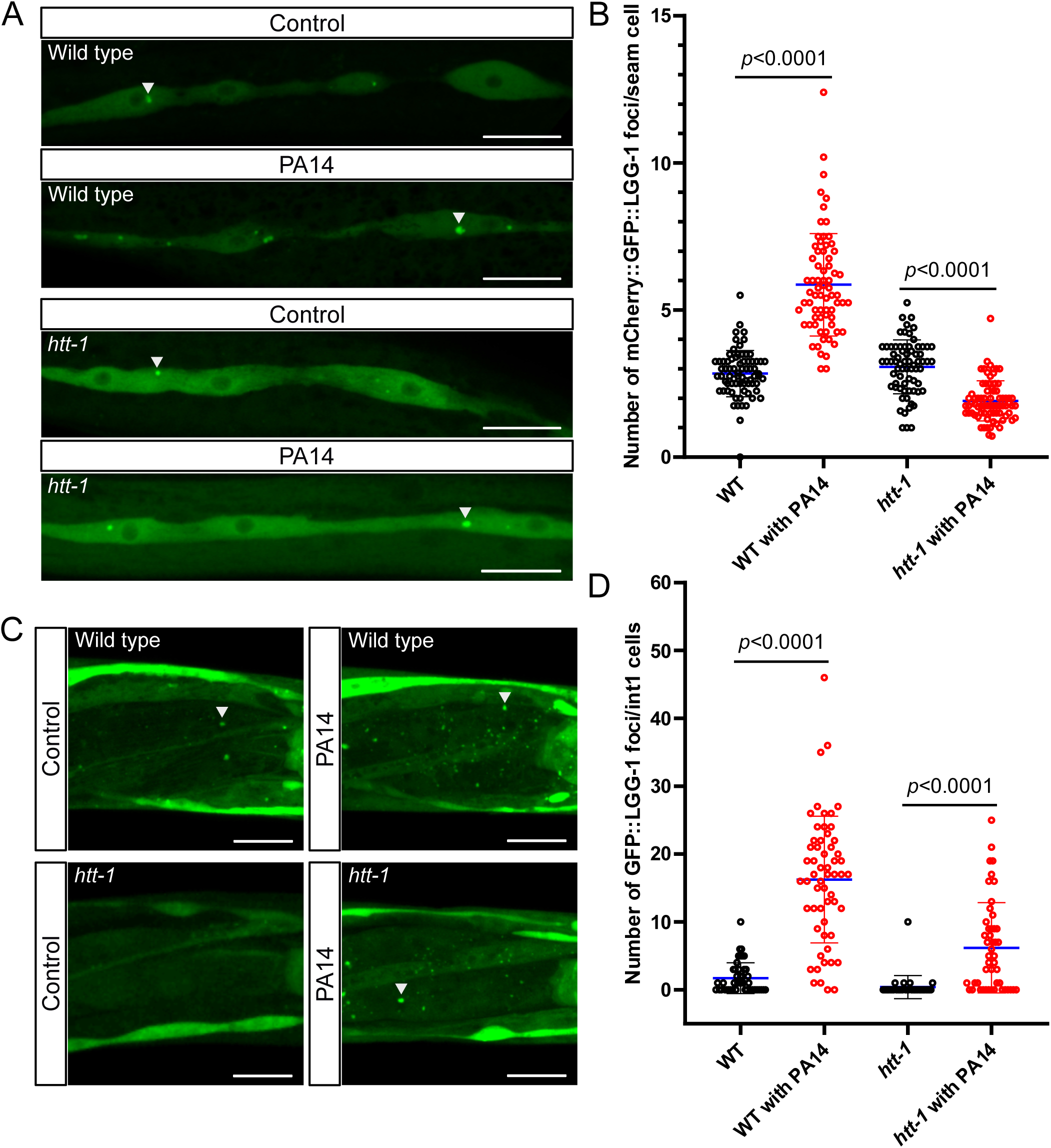
*htt-1* mediates robust autophagy induction in PA14-infected worms. (A) Confocal images of GFP::LGG-1 in the seam cells of wild-type (WT) and *htt-1(tm8121)* mutants. (B) Scatter plot showing the number of GFP::LGG-1 foci in infected (red dot) and uninfected (black dots) WT and *htt-1(tm8121)* mutants. Each dot represents the average number of GFP::LGG-1 foci counted across multiple seam cells in a single animal. (C) Confocal images of GFP::LGG-1 in int1 cells of WT and *htt-1(tm8121)* mutants. (D) Scatter plot showing the number of GFP::LGG-1 foci across all four int1 cells in a single infected (red dots) or uninfected (black dots) WT and *htt-1(tm8121)* mutant animals. Arrowheads indicate GFP::LGG-1 foci. Scale bars: 20 μm. *p* values are determined by unpaired t tests. Two-way ANOVA analyses are provided in S3 Table.

In wild-type N2 seam cells, the mean number of GFP::LGG-1 foci increased from 2.84 with OP50 to 5.86 with PA14 infection (*p* < 0.0001, Fig 5B and S3 Table). In contrast, *htt-1(tm8121)* seam cells exhibited a mean of 3.07 foci with OP50 and 1.91 foci with PA14 infection (*p* < 0.0001, Fig 5B and S3 Table). These results indicate that autophagy induction in seam cells is pronounced in wild-type N2, whereas no increase in autophagy activity is observed in *htt-1(tm8121)* mutants after infection. Similarly, in int1 cells, wild-type N2 exhibited a mean of 1.71 foci with OP50 and 16.24 foci with PA14 (*p* < 0.0001, Fig 5D and S3 Table), while *htt-1(tm8121)* mutants showed 0.4 foci with OP50 and 6.19 foci with PA14 (*p* < 0.0001, Fig 5D and S3 Table). Predicted least-squares mean from two-way ANOVA confirmed that the increase in autophagic activity, measured by GFP::LGG-1 puncta, was significantly greater in wild-type N2 compared to *htt-1(tm8121)* mutants upon PA14 infection (S3 Table). These findings indicate that the robust induction of autophagy observed in wild-type N2 is mediated by *htt-1,* whereas this response is diminished in *htt-1(tm8121)* mutants. Furthermore, autophagy activity in the anterior region was assessed, revealing that GFP::LGG-1 puncta were more prominently observed in areas associated with the pharynx, body wall muscle, hypodermis, nerve ring, and neurons of wild-type N2 compared to *htt-1(tm8121)* mutants (S5 Fig). Collectively, these findings suggest that the efficient induction of autophagy during PA14 infection requires the ERK pathway, CDC-48.2, and HTT-1, which together enhance survival in *C. elegans*.

Next, we aimed to investigate whether *htt-1* plays a role in responding to other stressors that also require autophagy as a defense mechanism. Thermal stress was chosen as a model. The heat shock response and autophagy are tightly linked. Various organisms, including *C. elegans,* induce autophagy under heat stress [26,27]. L4 stage animals were incubated at 34_ for 5 hours, and survival was monitored after 17 hours of incubation at 20_ (Fig 6A). Interestingly, *htt-1(tm8121)* mutants exhibited significantly reduced survival compared to wild-type N2 (*p* = 0.0257, Fig 6A). Expression of wild-type *htt-1* under its own promoter rescued the reduced survival after heat stress (N2 vs. *tm8121;*Ex[*htt-1p::htt-1*], *p* = 0.1917, Fig 6A). In addition, autophagy marker analysis revealed diminished autophagy activity in *htt-1(tm8121)* mutants relative to wild-type N2 after heat shock (Fig 6B and 6C). These findings suggest that *htt-1* is critical for robust induction of autophagy and survival under heat shock conditions. Moreover, the role of *htt-1* in the robust induction of autophagy and survival extends beyond immune defense, highlighting its potentially essential role in responding to other stressors that require autophagy.

**Fig 6.**
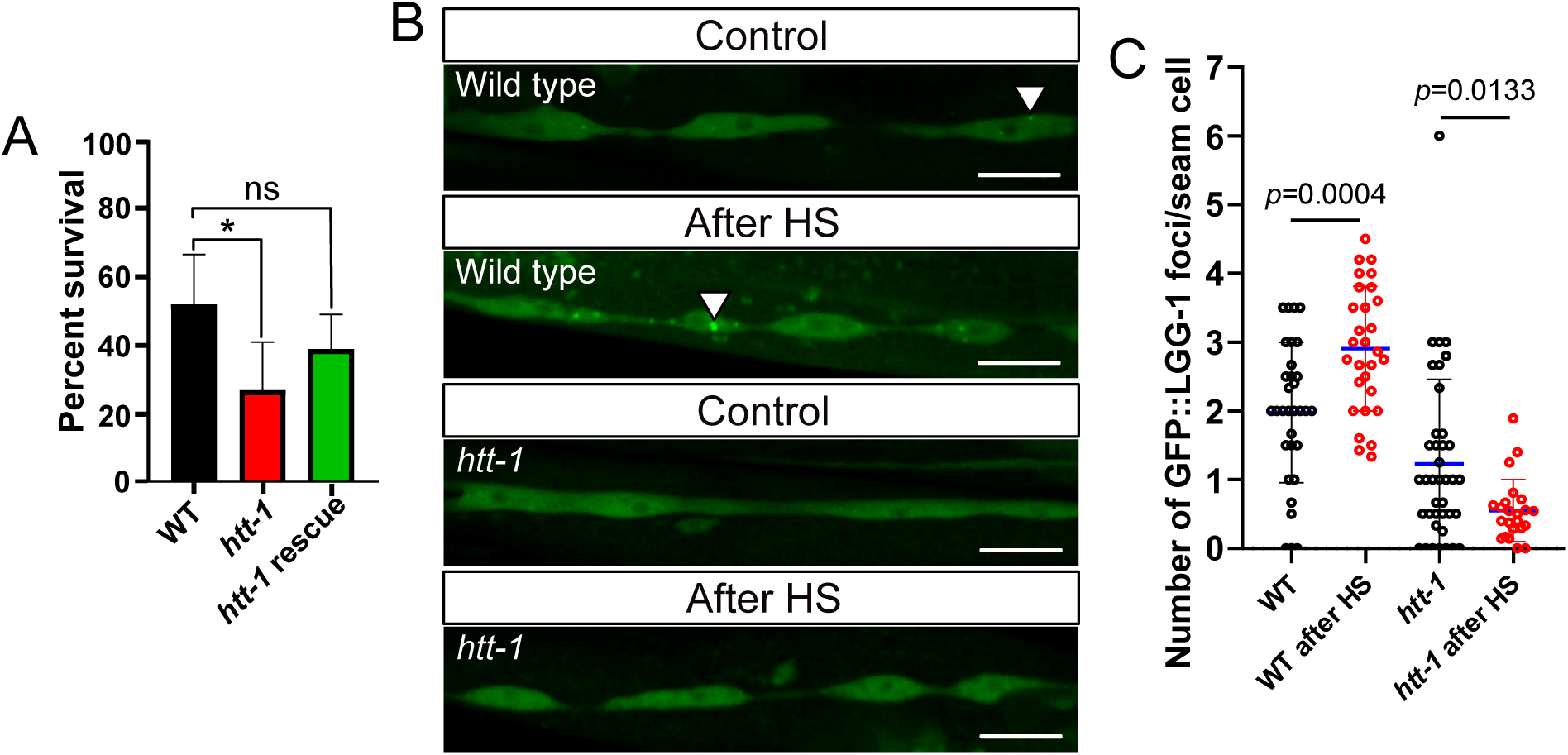
*htt-1* mediates robust autophagy induction in heat stressed animals. (A) Survival assessed in wild-type (WT) and *htt-1(tm8121)* following a 5-hour heat shock at 34_ and a 17-hour recovery period (25-35 animals per replicate, *n* = 2). ns (not significant) *p* > 0.05, * *p* ≤ 0.05, as determined by Unpaired t test. (B) Scatter plot showing the number of GFP::LGG-1 foci in heat-shocked (red dot) and the control (black dots) WT and *htt-1(tm8121)* mutants. Each dot represents the average number of GFP::LGG-1 foci counted across multiple seam cells in a single animal. *p* values are determined by unpaired t tests. Two-way ANOVA analyses are provided in S4 Table. Arrowheads indicate GFP::LGG-1 foci. Scale bars: 20 μm.

The expression of autophagy genes was monitored using standard quantitative PCR methods to further assess autophagy activity in wild-type and *htt-1* mutant animals [29]. Worms were exposed to pathogenic PA14 or non-pathogenic *Escherichia coli* OP50 bacteria for 12 hours at 25°C, followed by RT-PCR analysis. mRNA levels of autophagy genes, *atg-18, sqst-1,* and *lgg-1,* and the immune response lysozyme gene *lys-2* were measured. The *atg-18* gene encodes a homolog of the mammalian WIPI proteins (WD repeat protein interacting with phosphoinositides), which play a key role in the early stages of autophagy by facilitating phagophore formation [30-32]. SQST-1, a homolog of the mammalian p62/SQSTM1, acts as an autophagy adaptor, mediating cargo recognition and sequestration [33-35]. LGG-1, a homolog of the mammalian Atg8/LC3, is critical for autophagosome maturation and membrane elongation [36-39]. Additionally, LYS-2, which is primarily involved in immune responses, contributes to cellular cleanup by degrading bacterial and cellular debris [40-42].

The mRNA levels of *lgg-1, sqst-1,* and *atg-18* remained stable under PA14 infection compared OP50 control conditions in both wild-type N2 and *htt-1(tm8121)* mutants (*p* > 0.05, S6 Fig). While autophagy gene expression patterns were similar between wild-type N2 and *htt-1(tm8121)* mutants after 12 hours of infection, *htt-1(tm8121)* mutants exhibited a reduction in GFP::LGG-1 autophagosome markers in tissues near the head, as well as in int1 and seam cells along the body, compared to wild-type animals. This suggests that the decreased autophagy activity in *htt-1(tm8121)* mutants is not due to reduced expression of autophagy genes after 12 hours of infection.

Human HTT is a multifunctional protein, with nuclear-localized HTT involved in transcriptional regulation [1]. To investigate whether *C. elegans* HTT-1 also undergoes translocation, we analyzed the subcellular localization of HTT-1::GFP under standard laboratory conditions and stress conditions. HTT-1 remains cytoplasmic, both with and without 4 hours of heat shock at 35_ or 4 hours of PA14 infection. The cytoplasmic localization of HTT-1::GFP and stable autophagy related mRNA levels upon PA14 infection suggest that HTT-1’s regulation of robust autophagy response is not mediated by decreased transcript levels of autophagy genes in *htt-1(tm8121)* mutants compared to wild-type N2.

In contrast, *lys-2* mRNA levels were significantly elevated in both strains exposed to PA14 compared to OP50 controls (S6 Fig). In *htt-1(tm8121)* mutants, the increased *lys-2* expression during PA14 infection compared to uninfected controls may represent a compensatory response to reduced autophagy activity, potentially through enhanced lysosome-mediated degradation of immune-related debris. Despite the smaller *p* value for *htt-1(tm8121)* due to variability in the data, the increase in *lys-2* expression was greater compared to N2 (S5 Table).

## Discussion

The N-terminal of HTT is less conserved in lower eukaryotes. While the overall structure of HTT is conserved across species, the polyQ tract is poorly conserved in invertebrates. In humans, the polyQ region is followed by a proline-rich domain (PRD), a feature only found in mammals [43]. This region, however, plays a critical role in HD severity and progression. For example, the age of onset is inversely correlated with the length of the CAG repeat in the first exon of the human HTT gene. While HD typically manifests with late-onset symptoms, a juvenile form with earlier onset exists [44-46]. To model HD, many studies use transgenic animal expressing N-terminal fragments of HTT containing the polyQ tract, effectively demonstrating the toxicity of polyQ expansions. However, Htt is a big protein, and emerging evidence suggests that its central and C-terminal regions also harbor sequences essential for its diverse biological functions. Interestingly, deleting the normal polyQ stretch in HTT has demonstrated beneficial effects in HD mouse models. Expression of the polyQ-deleted form ameliorated disease phenotypes and extended lifespan, likely due to enhanced autophagy activity [47]. These findings highlight the functional importance of sequences throughout the entire HTT structure. Therefore, studying the normal functions of HTT utilizing various animal models is essential for gaining deeper insights into HD.

### Huntingtin’s role in selective autophagy

The relationship between huntingtin and autophagy has been previously studied. This study establishes a novel link between normal *htt-1* function, autophagy, and survival in *C. elegans.* Notably, while *C. elegans htt-1* mutation results in reduced autophagosome formation, HD models often show an increase in autophagosomes. In HD patients and mouse models, this autohpagy abnormality is characterized by elevated levels of autophagic vacuoles, the cargo adaptor p62, and the autophagosome marker LC3-II [48]. Despite increased autophagosome formation seeming to indicate enhanced autophagy, autophagy is, in fact, impaired in HD due to the toxic effects of polyQ-expanded mHTT. mHTT sequesters p62, limiting autophagosomal cargo recognition. These recognition defects often result in empty autophagosomes. Additionally, autophagy is defective in autophagosome transport, autolysosome formation, and cargo degradation. Collectively, these autophagic defects caused by mHTT contribute to neurotoxicity in HD [48].

In contrast, wild-type HTT has been shown to support autophagy through its role as a scaffold protein. The large, HEAT motif rich Htt acts as a scaffold protein essential for selective autophagy. The C-terminal domain of human Htt is structurally similar to yeast Atg11 and directly interacts with Atg11 interactors, which are essential for selective autophagy in yeast [49]. Additionally, studies using *Drosophila* and mammalian models have demonstrated that HTT functions in selective autophagy by modulating the activities of the cargo receptor p62 and the autophagy initiation kinase ULK1. HTT physically interacts with ULK1 to facilitate efficient selective autophagy. Interestingly, while HTT depletion does not impact starvation-induced activation of ULK1 kinase, it impairs ULK1 activation under proteotoxic stress. The selective regulation of ULK1 by HTT can be explained by its competition with the MTORC1 complex, another key regulator of ULK1. ULK1 interacts exclusively with either MTORC1 or HTT, forming the ULK1-HTT or ULK1-MTORC-1 complexes. The association between ULK1 and HTT is enhanced under stresses that induce selective autophagy, such as proteotoxicity, lipotoxicity, or mitophagy, but not starvation. This increased formation of ULK1-HTT complexes comes at the expense of MTOR, effectively freeing ULK1 from inhibition by MTORC1 and promoting selective autophagy. These findings suggest that HTT plays a role in the differential regulation of autophagosome biogenesis depending on basal or stress conditions [35,50].

HTT is crucial for selectivity in cargo recognition, which is crucial for autophagic quality control [35,50]. Similar to human HTT, *C. elegans* HTT-1’s role in autophagy may not extend to all types of stress. Interestingly, the loss of *htt-1* function was unable to fully suppress autophagy induction under heat shock or infection, indicating that while basal autophagy remains active, the activity of selective autophagy is reduced in *htt-1* mutant worms under stress.

During selective autophagy, HTT binds to autophagy proteins ULK1 and p62. HTT acts as a scaffold, bringing these proteins together to form a cargo-bound complex [35,50]. *C elegans* UNC-51 is the ortholog of mammalian ULK1 and ULK2. UNC-51 is known to function both as a primary initiator of autophagy and as a regulator of axon guidance, a function that is independent of autophagy [51-54]. Another binding protein, autophagy receptor p62/SQST-1, plays a key role in stress response and aging in *C. elegans* by facilitating the degradation of ubiquitinated cargo. Overexpression of *sqst-1* is sufficient to induce autophagy, extending lifespan and improving the fitness of proteostasis-defective worms. Additionally, *sqst-1* is required for autophagy induction upon heat shock, and the *sqst-1* mutation exhibits a phenotype similar to *htt-1* mutants [26]. Although suppressed autophagosome formation was observed in *htt-1* mutants along the body, the interactions between HTT-1 and the autophagy related proteins SQST-1 and UNC-51 are largely unknown. These potential similar interactors with HTT-1 and HTT would demonstrate robust functional conservation between the two orthologs, further revealing *htt-1*’s uncharacterized role as scaffolding in autophagy.

### *C. elegans* huntingtin ortholog, *htt-1,* in stress response

The ERK pathway, CDC-48.2 phosphorylation, and HTT-1’s coordinated activity in the intestine, along with autophagy regulation across various tissues are crucial for survival during infection [9] (Fig 7). In contrast, defects in ERK signaling, *htt-1*, and the autophagy cascade significantly reduces survival during infection. While intestine-specific *htt-1* expression rescues the *htt-1* mutant phenotype, *htt-1* may also function in other tissues to enhance survival during infection. While the intestine is the primary organ responsible for handling ingested stressors, the partial rescue observed with intestinal *htt-1* expression suggests that other tissues may still harbor autophagy-related defects due to the absence of normal *htt-1* function.

**Fig 7.**
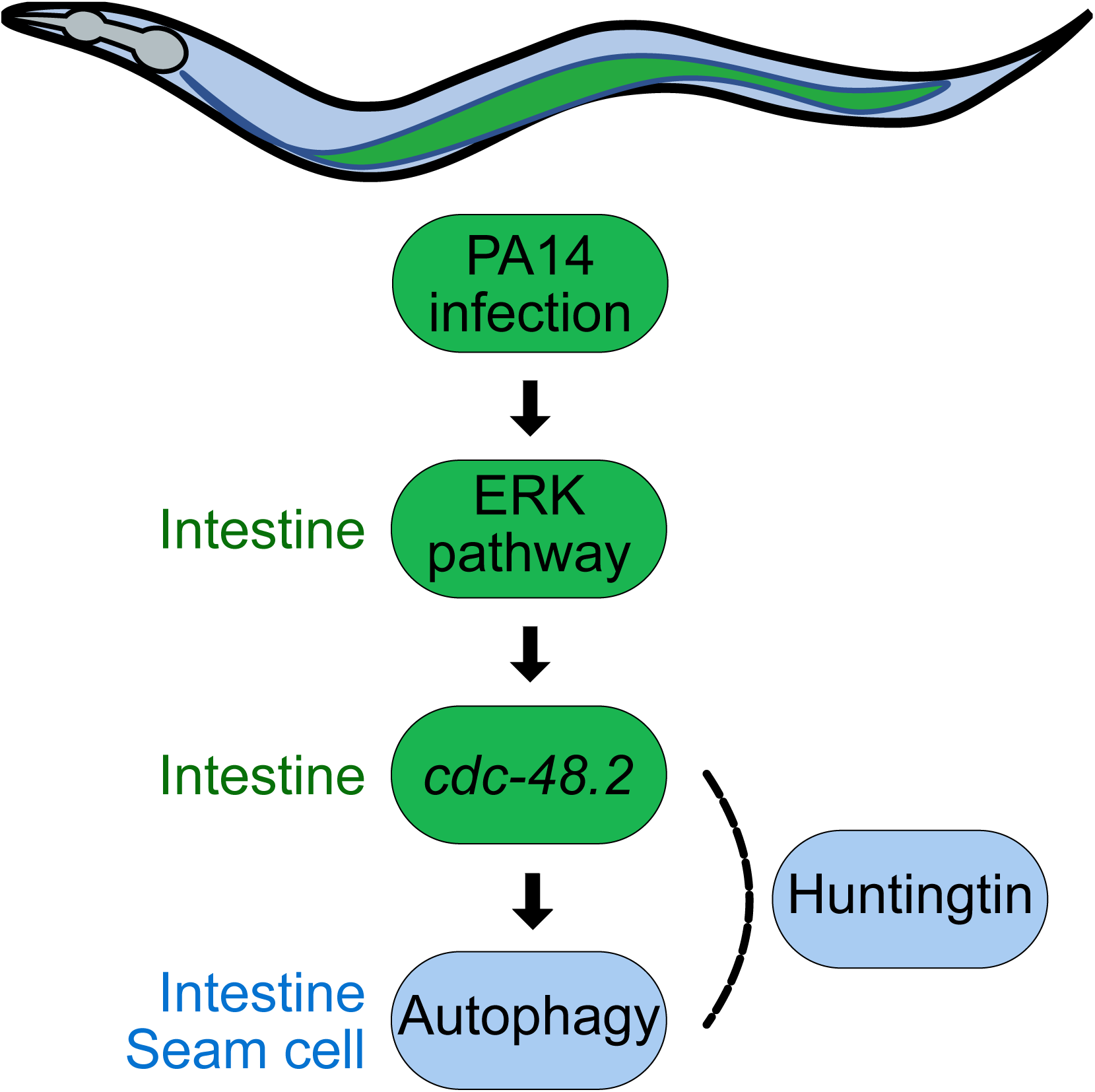
Model of *C. elegans* huntingtin *htt-1* regulation of stress response to PA14 infection in *C. elegans*.

## Materials and Methods

### *C. elegans* strains and maintenance

All *C. elegans* strains were grown under standard conditions at 20°C, as described by Brenner [55]. Strains used in this study include: Bristol N2, *let-60(n1046), lin-3(e1417),* MAH215: sqls11[*lgg-1p::mCherry::GFP::lgg-1 + rol-6*], *tm8121,* and *tm1959*. These strains were obtained from the Caenorhabditis Genetics Center and the National BioResource Project. Double mutants, *tm8121;let-60(n1046), tm8121;*sqls11[*lgg-1p::mCherry::GFP::lgg-1 + rol-6*], were generated by crossing. For transgenic strains with exogenous expression of *egl-3p::htt-1::gfp*, *htt-1p::htt-1::gfp*, *myo-3p::htt-1::gfp*, and *opt-2p::htt-1::gfp* vectors, or *htt-1p::HsHTT*(Q26 and Q76), the *unc-122p::RFP* vector was co-injected as a marker. Additionally, *egl-3p::mCherry*, *myo-3p:: mCherry,* and *opt-2p:: mCherry* vectors were co-injected with the *htt-1p::htt-1::gfp* vector.

### Molecular biology

The 3kb promoter region of *htt-1* with the full-length genomic DNA of *htt-1* (6193nt) and *gfp* were cloned into pPD117.01. For expression pattern analysis of *htt-1*, mCherry tagged promoters of *egl-3* (2000nt), *opt-2* (2012nt), and *myo-3* (2001nt) were subcloned into pPD117.01. For tissue specific expression of *htt-1*, promoters of *egl-3* (2000nt), *opt-2* (2012nt), and *myo-3* (2001nt) were subcloned into pPD117.01 containing *htt-1* (gDNA)*::gfp.* The 3 kb *htt-1* promoter and the full-length of HsHTT cDNA were cloned into pPD95.75.

### Measuring brood size

The average brood size was measured following the protocol described by Kwah and Jaramillo-Lambert [56]. Each L4 *C. elegans* was individually placed on a separate Nematode Growth Medium (NGM) plate and transferred to a new plate every one or two days. After removing the worms, the number of hatched progenies were counted for every strain. The obtained data were analyzed using an unpaired t test (GraphPad Prism).

### Measuring lifespan

Lifespan was measured using the modified protocol described by Sutphin and Kaeberlein [57]. Day 1 adult *C. elegans* were allowed to lay eggs on a 50 mm plate at 20℃ for 2 hours. After the progeny reached the L4 stage, 25-30 animals were transferred to each plate. Survival was monitored by counting live and dead animals every one or two days. Survival rate data were analyzed using survival curve comparison (GraphPad Prism).

### *C. elegans* survival assay under *Pseudomonas aeruginosa* PA14 infection

Slow-killing assay was performed by modifying the protocol described by Tan [58]. To assess *C. elegans* survival under *P. aeruginosa* PA14 infection, pathogen-seeded slow killing plates were prepared by growing PA14 overnight in LB medium at 37°C. A 10uL of the culture was seeded and spread onto the center of 50 mm NGM plates, which were subsequently incubated at 37°C overnight to allow bacterial colonization. Approximately 30 synchronized *C. elegans* at the L4 stage were transferred to these PA14-seeded plates and incubated at 25°C. Survival rates were measured every 8-12 hours, starting 24 hours post-exposure, and continued until all worms were dead or censored. To investigate the effects of U0126 treatment on *P. aeruginosa* PA14 infection, *C. elegans* were exposed *P. aeruginosa* PA14-seeded plates containing 8 μM U0126 (Sigma-Aldrich U0126 ethanolate), with survival recorded as described. Additionally, RNAi-treated L4 stage *C. elegans* were transferred to PA14-seeded plates to assess the impact of gene knockdown on survival during infection. Survival rate data were analyzed using survival curve comparison (GraphPad Prism).

### Measuring lawn occupancy

Lawn occupancy was assessed following the protocol described by Singh and Aballay [11]. 20μL of overnight grown *P. aeruginosa* PA14 was seeded on the center of a 50mm slow-killing assay plate and incubated for 12 hours. Approximately 25 *C. elegans* were placed at the edges of the plate, outside the bacterial lawn. The number of animals inside and outside the lawn was recorded, and lawn occupancy was calculated as N_on_ lawn/N_total_.

### RNA interference

RNAi bacterial clones were obtained from the J. Ahringer library. For the *mek-2* RNAi, the *mek-2* cDNA (bases 143-1870) was cloned into the L4440 vector. Control (L4440) or target RNAi clones were introduced into *E. coli* HT115. L4 worms were transferred to plates seeded with RNAi-feeding bacteria and maintained for one day. They were then removed, and their progeny reared on RNAi-feeding bacteria were used for subsequent experiments.

### Autophagy activity analysis of *C. elegans* exposed to PA14 or heat shock

For infection experiments, L4-staged sqls11[*lgg-1p::mCherry::GFP::lgg-1 + rol-6*] animals, either with control N2 or mutant *tm8121* backgrounds, were transferred to plates seeded with either *E. coli* OP50 or *P. aeruginosa* PA14. The worms were incubated at 25°C for 12 hours. For heat shock experiments, L4-staged animals were subjected to a 2-hour heat shock at 37°C, followed by a recovery period of 30 minutes to 1 hour at 20°C. After treatment, the worms were anesthetized in M9 medium containing 0.1% sodium azide and mounted for imaging. GFP::LGG-1 puncta were quantified from fluorescence images acquired using a confocal microscope (ZEISS LSM700; Carl Zeiss) and analyzed with ZEN software (Carl Zeiss). Quantification focused on the first four intestinal cells (int1) and more than three seam cells per worm along the z-stack.

### Quantitative RT-PCR

*E. coli* OP50 or *P. aeruginosa* PA14 seeded 50mm NGM plates were prepared. About 50 L4-stage synchronized N2 and *htt-1(tm8121)* animals were transferred to six plates for each strain for each condition. After a 12-hour incubation at 25℃, all worms were collected with M9 medium using mouth pipette. Total RNA was isolated using TRI reagent (Molecular Research Center) by the freeze-thaw method. cDNA was synthesized with Omniscript RT Kit (Qiagen) using oligo(dT) primer. Quantitative real-time PCR was performed using Bio-Rad iQ SYBR Green supermix in Bio-Rad CFX connect as described in manufacturer’s manual. For sequences of primers used for quantitative RT-PCR are listed in S6 Table. mRNA levels of target genes were normalized to the mean of *act-1/3* housekeeping gene. Data are displayed as relative values compared with controls. Data were analyzed using unpaired t test (GraphPad Prism).

### Subcellular localization of HTT-1::GFP in *C. elegans*

Approximately 20 of L4-staged *htt-1p::htt-1::gfp*-expressing animals were subjected to described experimental conditions. Animals were either exposed to *P. aeruginosa* PA14 or *E. coli* OP50 for 4 hours at 25°C or treated with a 1-hour heat shock at 35℃ followed by a 1-hour recovery at 20℃. After treatment, the animals were immobilized with 2.5 mM levamisole, and HTT-1::GFP subcellular localization was observed using a confocal microscope (ZEISS LSM700; Carl Zeiss) and analyzed with ZEN software (Carl Zeiss).

### Measuring survival after heat shock

Survival after heat shock was evaluated using a modified version of the protocol described by Zevian and Yanowitz [59]. Approximately 30 synchronized L4-stage *C. elegans* were placed on each plate, with three replicate plates per strain. The plates were incubated at 34℃ for 5 hours to induce heat stress. Following the treatment, the worms were maintained at 20℃, and survival was assessed 17 hours later.

## Acknowledgments

We thank J. Ahringer and M. Vidal for the RNAi plasmids. Additionally, we acknowledge the Caenorhabditis Genetic Center and the National BioResource Project for providing the strains.

## Notes

### Competing Interest Statement

The authors have declared no competing interest.

### Summary of Updates

We have updated all data and figures to make the manuscript understandable.

