## Supplementary figures and tables for "*C. elegans* huntingtin, *htt-1,* promotes robust autophagy induction and survival under stress conditions"

### Supporting information

**S1 Table. Statistical analyses.** The number represents the total number of animals tested. *p* values are calculated relative to the wild-type control unless otherwise specified.

Figure 1C

| Genotype | Mean Lifespan (days) | Number | *P* value | *P* value summary |
|---|---|---|---|---|
| Wild type | 17 | 166 | | |
| *htt-1(tm8121)* | 16 | 163 | 0.0981 | ns |
| *htt-1(tm1959)* | 17 | 141 | 0.5181 | ns |

Figure 1D

| Genotype | Mean | Number | *P* value | *P* value summary |
|---|---|---|---|---|
| Wild type | 299.2 | 15 | | |
| *htt-1(tm8121)* | 285.8 | 16 | 0.6668 | ns |
| *htt-1(tm1959)* | 293.2 | 12 | 0.1316 | ns |

Figure 2A

| Genotype | Median Lifespan (hours) | Number | *P* value | *P* value summary |
|---|---|---|---|---|
| Wild type | 60 | 259 | | |
| *htt-1(tm8121)* | 48 | 277 | <0.0001 | **** |
| *htt-1(tm1959)* | 48 | 280 | <0.0001 | **** |

Figure 2B

| Genotype | Median Lifespan (hours) | Number | *P* value | *P* value summary |
|---|---|---|---|---|
| Wild type | 45 | 321 | | |
| *htt-1(tm8121)* | 45 | 391 | <0.0001 | **** |
| *tm8121*;Ex[*htt-1p::htt-1::gfp*] | 45 | 381 | 0.0859 | ns |

Figure 2C

| Genotype | Median Lifespan (hours) | Number | *P* value | *P* value summary |
|---|---|---|---|---|
| Wild type | 48 | 168 | | |
| *htt-1(tm8121)* | 36 | 196 | <0.0001 | **** |

Figure 2D

| Genotype | Median Lifespan (hours) | Number | *P* value | *P* value summary |
|---|---|---|---|---|
| Wild type | 36 | 120 | | |
| *htt-1(tm8121)* | 24 | 100 | 0.0003 | *** |

Figure 2E

| Genotype | Median Lifespan (hours) | Number | *P* value | *P* value summary |
|---|---|---|---|---|
| Wild type | 52 | 256 | | |
| *htt-1(tm8121)* | 48 | 257 | <0.0001 | **** |
| *tm8121*;Ex[intestine *htt-1::gfp*] | 48 | 218 | 0.0003 | *** |
| *tm8121*;Ex[Neuron *htt-1::gfp*] | 48 | 246 | <0.0001 | **** |
| *tm8121*;Ex[Muscle *htt-1::gfp*] | 48 | 262 | <0.0001 | **** |

| Genotype | *P* value | *P* value summary |
|---|---|---|
| *tm8121* vs *tm8121*;Ex[intestine *htt-1::gfp*] | <0.0001 | **** |
| *tm8121* vs *tm8121*;Ex[Neuron *htt-1::gfp*] | 0.8986 | ns |
| *tm8121* vs *tm8121*;Ex[Muscle *htt-1::gfp*] | 0.6157 | ns |

Figure 3

| Genotype | Median Lifespan (hours) | Number | *P* value | *P* value summary |
|---|---|---|---|---|
| Wild type | 40 | 202 | | |
| *htt-1(tm8121)* | 40 | 257 | <0.0001 | **** |
| *tm8121*;Ex[*htt-1p::HsHTT(Q26)*] | 40 | 250 | 0.3619 | ns |
| *tm8121*;Ex[*htt-1p::mHsHTT(Q76)*] | 32.5 | 285 | <0.0001 | **** |

| Genotype | *P* value | *P* value summary |
|---|---|---|
| *tm8121* vs *tm8121*;Ex[*htt-1p::HsHTT(Q26)*] | <0.0001 | **** |
| *tm8121* vs *tm8121*;Ex[*htt-1p::HsHTT(Q76)*] | 0.0009 | *** |

Figure 4A

| Genotype | Median Lifespan (hours) | Number | *P* value | *P* value summary |
|---|---|---|---|---|
| Wild type | 53 | 225 | | |
| N2 *lin-3* (RNAi) | 53 | 246 | <0.0001 | **** |
| *htt-1(tm8121)* | 45 | 298 | <0.0001 | **** |
| *tm8121 lin-3* (RNAi) | 45 | 270 | <0.0001 | **** |

| Genotype | *P* value | *P* value summary |
|---|---|---|
| *tm8121* vs *tm8121 lin-3* (RNAi) | 0.8196 | ns |

Figure 4B

| Genotype | Median Lifespan (hours) | Number | *P* value | *P* value summary |
|---|---|---|---|---|
| Wild type | 50 | 210 | | |
| N2 *let-60* (RNAi) | 50 | 208 | <0.0001 | **** |
| *htt-1(tm8121)* | 40 | 229 | <0.0001 | **** |
| *tm8121 let-60* (RNAi) | 50 | 274 | <0.0001 | **** |

| Genotype | *P* value | *P* value summary |
|---|---|---|
| *tm8121* vs *tm8121 let-60* (RNAi) | 0.1506 | ns |

Figure 4C

| Genotype | Median Lifespan (hours) | Number | *P* value | *P* value summary |
|---|---|---|---|---|
| Wild type | 50 | 248 | | |
| N2 *mek-2* (RNAi) | 50 | 137 | 0.0025 | ** |
| *htt-1(tm8121)* | 40 | 296 | <0.0001 | **** |
| *tm8121 mek-2* (RNAi) | 40 | 176 | <0.0001 | **** |

| Genotype | *P* value | *P* value summary |
|---|---|---|
| *tm8121* vs *tm8121 mek-2* (RNAi) | 0.7912 | ns |

Figure 4D

| Genotype | Median Lifespan (hours) | Number | *P* value | *P* value summary |
|---|---|---|---|---|
| Wild type | 60 | 262 | | |
| N2 *mpk-1* (RNAi) | 45 | 258 | <0.0001 | **** |
| *htt-1(tm8121)* | 45 | 265 | <0.0001 | **** |
| *tm8121 mpk-1* (RNAi) | 45 | 263 | <0.0001 | **** |

| Genotype | *P* value | *P* value summary |
|---|---|---|
| *tm8121* vs *tm8121 mpk-1* (RNAi) | 0.1898 | ns |

Figure 4E

| Genotype | Median Lifespan (hours) | Number | *P* value | *P* value summary |
|---|---|---|---|---|
| Wild type | 60 | 326 | | |
| *lin-3(e1417)* | 45 | 279 | <0.0001 | **** |
| *htt-1(tm8121)* | 45 | 277 | <0.0001 | **** |
| *htt-1(tm8121);lin-3(e1417)* | 45 | 228 | <0.0001 | **** |

| Genotype | *P* value | *P* value summary |
|---|---|---|
| *tm8121* vs *htt-1(tm8121);lin-3(e1417)* | 0.3982 | ns |

Figure 4F

| Genotype | Median Lifespan (hours) | Number | *P* value | *P* value summary |
|---|---|---|---|---|
| Wild type | 60 | 185 | | |
| N2 with U0126 | 45 | 189 | <0.0001 | **** |
| *htt-1(tm8121)* | 45 | 252 | <0.0001 | **** |
| *tm8121* with *U0126* | 45 | 321 | <0.0001 | **** |

| Genotype | *P* value | *P* value summary |
|---|---|---|
| *tm8121* vs *tm8121* with *U0126* | 0.3017 | ns |

Figure 4G

| Genotype | Median Lifespan (hours) | Number | *P* value | *P* value summary |
|---|---|---|---|---|
| Wild type | 55 | 165 | | |
| *let-60(n1046)* | 55 | 470 | 0.0011 | ** |
| *htt-1(tm8121)* | 48 | 371 | <0.0001 | **** |
| *htt-1(tm8121);let-60(n1046)* | 48 | 234 | <0.0001 | **** |

| Genotype | *P* value | *P* value summary |
|---|---|---|
| *tm8121* vs *htt-1(tm8121);let-60(n1046)* | 0.4936 | ns |

Figure 4H

| Genotype | Median Lifespan (hours) | Number | *P* value | *P* value summary |
|---|---|---|---|---|
| Wild type | 50 | 242 | | |
| N2 *cdc-48.2* (RNAi) | 50 | 250 | 0.0002 | *** |
| *htt-1(tm8121)* | 50 | 275 | <0.0001 | **** |
| *tm8121 cdc-48.2 (RNAi)* | 50 | 280 | <0.0001 | **** |

| Genotype | *P* value | *P* value summary |
|---|---|---|
| *tm8121* vs *tm8121 cdc-48.2 (RNAi)* | 0.7361 | ns |

Figure 4I

| Genotype | Median Lifespan (hours) | Number | *P* value | *P* value summary |
|---|---|---|---|---|
| Wild type | 50 | 318 | | |
| N2 *bec-1* (RNAi) | 50 | 336 | 0.0003 | *** |
| *htt-1(tm8121)* | 45 | 414 | <0.0001 | **** |
| *tm8121 bec-1 (RNAi)* | 45 | 309 | <0.0001 | **** |

| Genotype | *P* value | *P* value summary |
|---|---|---|
| *tm8121* vs *tm8121 cdc-48.2 (RNAi)* | 0.5008 | ns |

Fig 6A

| Genotype | Number | *P* value | *P* value summary |
|---|---|---|---|
| Wild type | 195 | | |
| *htt-1(tm8121)* | 256 | 0.0257 | * |
| *tm8121*;Ex[*htt-1p::htt-1::gfp*] | 171 | 0.1917 | ns |

S3A Fig

| Genotype | Median Lifespan (hours) | Number | *P* value | *P* value summary |
|---|---|---|---|---|
| Wild type | 55 | 224 | | |
| N2 *clp-1*(RNAi) | 50 | 214 | 0.8282 | ns |
| *htt-1(tm8121)* | 45 | 309 | <0.0001 | **** |
| *tm8121 clp-1 (*RNAi) | 45 | 287 | <0.0001 | **** |

| Genotype | *P* value | *P* value summary |
|---|---|---|
| *tm8121* vs *tm8121 clp-1* (RNAi) | 0.7906 | ns |

S3B Fig

| Genotype | Median Lifespan (hours) | Number | *P* value | *P* value summary |
|---|---|---|---|---|
| Wild type | 65 | 174 | | |
| N2 *ced-3* (RNAi) | 65 | 162 | 0.2043 | ns |
| *htt-1(tm8121)* | 54 | 175 | <0.0001 | **** |
| *tm8121 ced-3 (*RNAi) | 54 | 194 | <0.0001 | **** |

| Genotype | *P* value | *P* value summary |
|---|---|---|
| *tm8121* vs *tm8121 ced-3* (RNAi) | 0.157 | ns |

S4A Fig

| Genotype | Median Lifespan (hours) | Number | *P* value | *P* value summary |
|---|---|---|---|---|
| Wild type | 50 | 228 | | |
| N2 *daf-16* (RNAi) | 50 | 212 | 0.7156 | ns |
| *htt-1(tm8121)* | 40 | 255 | <0.0001 | **** |
| *tm8121 daf-16 (*RNAi) | 40 | 275 | <0.0001 | **** |

| Genotype | *P* value | *P* value summary |
|---|---|---|
| *tm8121* vs *tm8121 daf-16* (RNAi) | <0.0001 | **** |

S4B Fig

| Genotype | Median Lifespan (hours) | Number | *P* value | *P* value summary |
|---|---|---|---|---|
| Wild type | 47 | 310 | | |
| N2 *dkf-2* (RNAi) | 47 | 265 | <0.0001 | **** |
| *htt-1(tm8121)* | 47 | 254 | <0.0001 | **** |
| *tm8121 dkf-2 (*RNAi) | 38 | 263 | <0.0001 | **** |

| Genotype | *P* value | *P* value summary |
|---|---|---|
| *tm8121* vs *tm8121 dkf-2* (RNAi) | <0.0001 | **** |

S4C Fig

| Genotype | Median Lifespan (hours) | Number | *P* value | *P* value summary |
|---|---|---|---|---|
| Wild type | 45 | 443 | | |
| N2 *egl-30* (RNAi) | 35 | 366 | <0.0001 | **** |
| *htt-1(tm8121)* | 45 | 335 | <0.0001 | **** |
| *tm8121 egl-30* (RNAi) | 35 | 342 | <0.0001 | **** |

| Genotype | *P* value | *P* value summary |
|---|---|---|
| *tm8121* vs *tm8121 egl-30* (RNAi) | <0.0001 | **** |

S4D Fig

| Genotype | Median Lifespan (hours) | Number | *P* value | *P* value summary |
|---|---|---|---|---|
| Wild type | 50 | 296 | | |
| N2 *fshr-1* (RNAi) | 40 | 138 | <0.0001 | **** |
| *htt-1(tm8121)* | 40 | 318 | <0.0001 | **** |
| *tm8121 fshr-1* (RNAi) | 40 | 168 | <0.0001 | **** |

| Genotype | *P* value | *P* value summary |
|---|---|---|
| *tm8121* vs *tm8121 fshr-1* (RNAi) | <0.0001 | **** |

S4E Fig

| Genotype | Median Lifespan (hours) | Number | *P* value | *P* value summary |
|---|---|---|---|---|
| Wild type | 50 | 166 | | |
| N2 *pmk-1* (RNAi) | 40 | 214 | <0.0001 | **** |
| *htt-1(tm8121)* | 40 | 195 | <0.0001 | **** |
| *tm8121 pmk-1* (RNAi) | 30 | 246 | <0.0001 | **** |
| Genotype | | | *P* value | *P* value summary |
| *tm8121* vs *tm8121 pmk-1* (RNAi) | | | <0.0001 | **** |

**S2 Table. Statistical analyses of PA14 lawn occupancy.**

Multiple t test and two-way AVOVA test results.

| Time(h) | Discovery? | P value | Mean of N2 | Mean of tm8121 | Difference | SE of difference | t ratio | df | q value |
|---|---|---|---|---|---|---|---|---|---|
| 2 | No | 0.39032 | 0.7702 | 0.8307 | -0.06055 | 0.0683 | 0.8865 | 14 | 0.315378 |
| 4 | No | 0.126908 | 0.8135 | 0.8922 | -0.07871 | 0.0485 | 1.623 | 14 | 0.128177 |
| 8 | No | 0.92615 | 0.659 | 0.6458 | 0.01322 | 0.1404 | 0.09416 | 16 | 0.623608 |
| 12 | Yes | 0.000163 | 0.4505 | 0.1776 | 0.273 | 0.05582 | 4.89 | 16 | 0.00066 |
| 16 | Yes | 0.005339 | 0.2613 | 0.1172 | 0.1441 | 0.04475 | 3.221 | 16 | 0.00719 |
| 24 | Yes | 0.003065 | 0.1271 | 0.02921 | 0.09791 | 0.0281 | 3.484 | 16 | 0.006191 |

| 2way ANOVA anlysis of lawn occupancy | |
|---|---|
| Difference between row means | |
| Predicted (LS) mean of N2 | 0.5136 |
| Predicted (LS) mean of tm8121 | 0.4488 |
| Difference between predicted means | 0.06483 |
| SE of difference | 0.03063 |
| 95% CI of difference | 0.003991 to 0.1257 |

**S3 Table. Statistical analyses of GFP::LGG-1 foci measurements in PA14-infected animals.**

Th N represents the total number of animals tested. Results from two-way ANOVA are shown.

Fig 5B. GFP::LGG-1 foci in seam cells

| Bacteria | OP50 | | | PA14 | | |
|---|---|---|---|---|---|---|
| Strain | Mean | SD | N | Mean | SD | N |
| N2 | 2.8385618 | 0.7863189 | 74 | 5.8594558 | 1.7423243 | 70 |
| tm8121 | 3.0705045 | 0.9133753 | 70 | 1.9115238 | 0.6828054 | 75 |

| 2way ANOVA anlysis of LGG-1 foci in seam cells | |
|---|---|
| Difference between row means | |
| Predicted (LS) mean of N2 | 4.349 |
| Predicted (LS) mean of tm8121 | 2.491 |
| Difference between predicted means | 1.858 |
| SE of difference | 0.1298 |
| 95% CI of difference | 1.602 to 2.114 |

Fig 5D. GFP::LGG-1 foci in int1 cells

| Bacteria | OP50 | | | PA14 | | |
|---|---|---|---|---|---|---|
| Strain | Mean | SD | N | Mean | SD | N |
| N2 | 1.7111111 | 2.2625631 | 45 | 16.241379 | 9.3270223 | 58 |
| tm8121 | 0.4 | 1.7012106 | 35 | 6.1875 | 6.6577622 | 48 |

| 2way ANOVA anlysis of LGG-1 foci in int1 cells | |
|---|---|
| Difference between row means | |
| Predicted (LS) mean of N2 | 8.976 |
| Predicted (LS) mean of tm8121 | 3.294 |
| Difference between predicted means | 5.682 |
| SE of difference | 0.9482 |
| 95% CI of difference | 3.812 to 7.553 |

**S4 Table. Statistical analyses of GFP::LGG-1 foci measurements in seam cells of heat-stressed animals.**

N represents the total number of animals tested. Results from two-way ANOVA are shown.

Fig 6B

| Bacteria | OP50 | | | PA14 | | |
|---|---|---|---|---|---|---|
| Strain | Mean | SD | N | Mean | SD | N |
| N2 | 1.976 | 1.021 | 32 | 2.904 | 0.9063 | 29 |
| tm8121 | 1.23 | 1.227 | 39 | 0.55 | 0.4504 | 23 |

| 2way ANOVA anlysis of LGG-1 foci in seam cells | |
|---|---|
| Difference between row means | |
| Predicted (LS) mean of N2 | 2.44 |
| Predicted (LS) mean of tm8121 | 0.89 |
| Difference between predicted means | 1.55 |
| SE of difference | 0.1821 |
| 95% CI of difference | 1.190 to 1.911 |

**S5 Table. Statistical analyses of RT-PCR experiments.**

Unpaired t test and two-way ANOVA test results.

| Table analyzed | | Mean | *P* value | *P* value summary |
|---|---|---|---|---|
| *lgg-1* | N2<br>N2 with PA14 | 1<br>1.296 | 0.4165 | ns |
| *lgg-1* | tm8121<br>tm8121 with PA14 | 1<br>0.7649 | 0.7197 | ns |
| *sqst-1* | N2<br>N2 with PA14 | 1<br>1.163 | 0.5919 | ns |
| *sqst-1* | tm8121<br>tm8121 with PA14 | 1<br>1.523 | 0.5104 | ns |
| *atg-18* | N2<br>N2 with PA14 | 1<br>1.472 | 0.629 | ns |
| *atg-18* | tm8121<br>tm8121 with PA14 | 1<br>7.469 | 0.1417 | ns |
| *lys-2* | N2<br>N2 with PA14 | 1<br>3.817 | 0.0022 | ** |
| *lys-2* | tm8121<br>tm8121 with PA14 | 1<br>22.34 | 0.0238 | * |

| 2way ANOVA anlysis of *lys-2* expression levels | |
|---|---|
| Difference between row means | |
| Predicted (LS) mean of N2 | 2.409 |
| Predicted (LS) mean of tm8121 | 11.67 |
| Difference between predicted means | -9.259 |
| SE of difference | 3.011 |
| 95% CI of difference | -16.20 to -2.315 |

**S6 Table. Sequence of quantitative RT-PCR primers used in this study.**

| Gene | Primer sequence 5' → 3' | |
|---|---|---|
| Experimental genes | | |
| *act-1/3* | Fwd | ACG CCA ACA CTG TTC TTT CC |
| | Rev | GAT GAT CTT GAT CTT CAT GGT TGA |
| *atg-18* | Fwd | ACA ACA AGC CAG AAG CGT CT |
| | Rev | CTG GTT TTT ATG TGA AAC AAG TG |
| *lgg-1* | Fwd | GCT CCA TGA CTT GGA TAA GAA |
| | Rev | CGT GAT GGT CCT GGT AGA G |
| *sqst-1* | Fwd | CGA GAG CCT CAG TCC ATC A |
| | Rev | GAC GAT CGA ATG TCT TTT CAG |
| *lys-2* | Fwd | CCG TGG ATT TGT TCC AAC TG |
| | Rev | GGG GAA GTAACC TGA ATC CA |

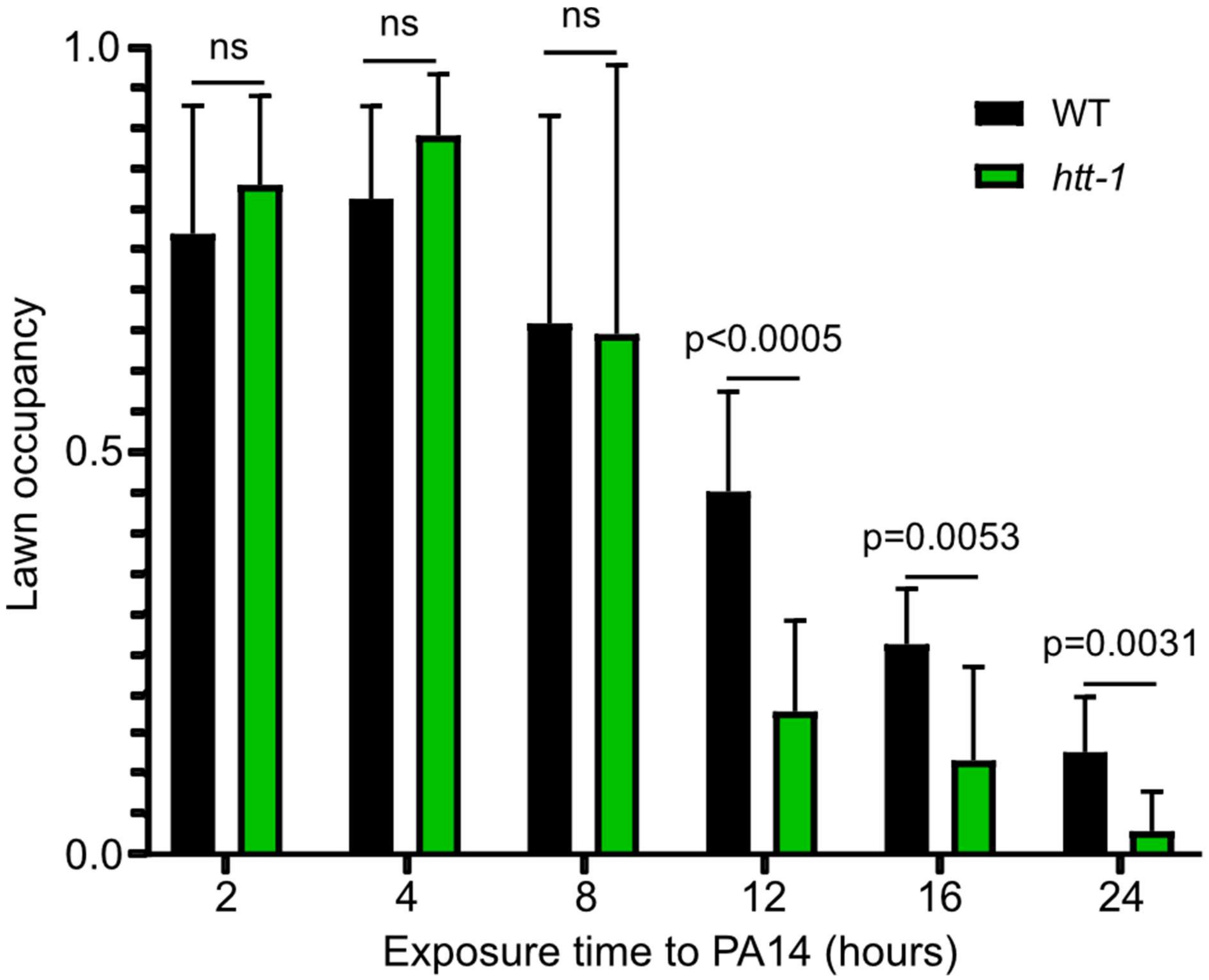


**S1 Fig. Wild-type and *htt-1* mutants exhibit similar pathogen avoidance behavior during the initial stages of exposure to PA14.** Lawn occupancy of animals on PA14 was evaluated over 24 hours in wild-type N2 and *htt-1(tm8121)* mutant worms. (70-90 animals per replicate, $n = 3$). ns (not significant) $p > 0.05$, * $p \leq 0.05$, ** $p \leq 0.01$, *** $p \leq 0.001$, **** $p \leq 0.0001$ as determined by Multiple t tests. Two-way ANOVA statistical analyses are provided in the S2 Table. Error bars represent SD.

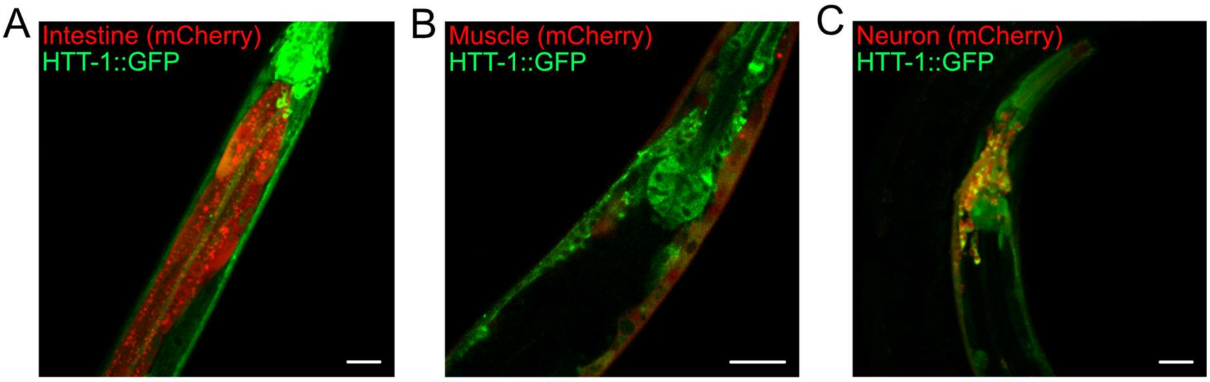


**S2 Fig. HTT-1 is expressed in multiple tissues, including the intestine, muscle, and neurons.** (A) Confocal image showing the expression of *htt-1::gfp* and the intestine marker *opt-2p::mCherry* expression in wild-type (WT) animals. (B) Confocal image showing *htt-1::gfp* and the muscle marker *myo-3p::mCherry* expression in WT animals. (C) Confocal image showing *htt-1::gfp* and the pan-neuronal marker *egl-3p::mCherry* expression in WT animals. Scale bars: 20 µm.

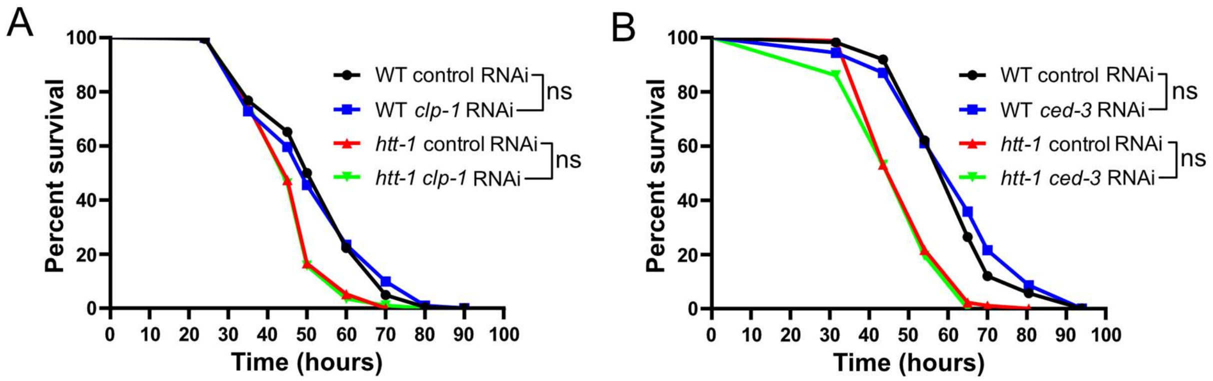


**S3 Fig. Reduced survival of *htt-1* mutants during PA14 infection is not linked to increased necrosis or apoptosis.** (A) Percent survival of wild-type (WT) and *htt-1(tm8121)* animals with empty vector L4440 or *clp-1* RNAi during PA14 infection (25-35 animals per replicate, $n = 3$). (B) Percent survival of WT and *htt-1(tm8121)* animals with empty vector L4440 or *ced-3* RNAi during PA14 infection (25-35 animals per replicate, $n = 2$). ns (not significant) $p > 0.05$ and **** $p \leq 0.0001$ as determined by Log rank test. Detailed statistical analyses are provided in the S1 Table.

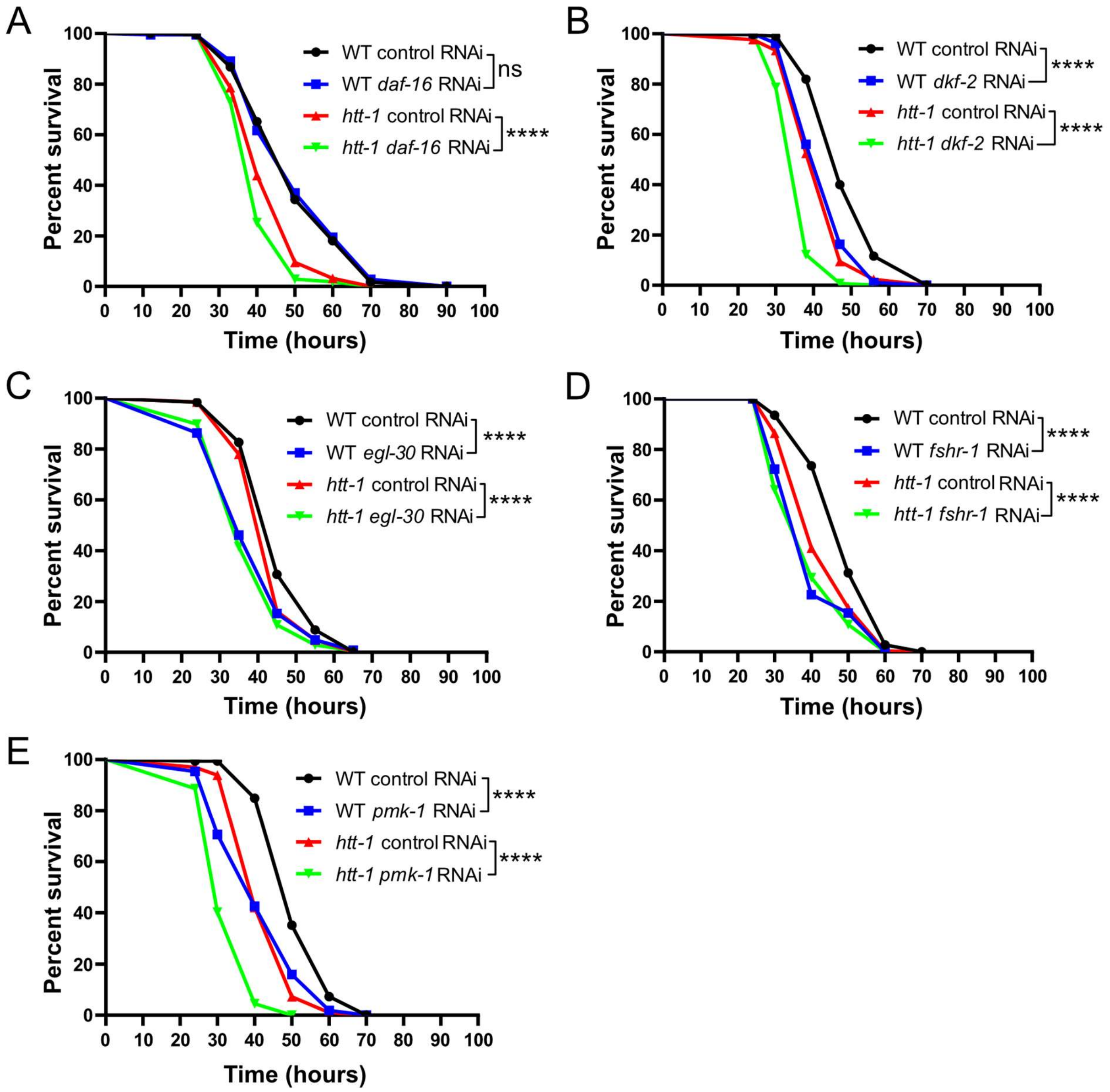


**S4 Fig. *htt-1* functions independently of the DAF-16, DKF-2, EGL-30, FSHR-1, and PMK-1 pathways to promote survival during PA14 infection.**

**S4 Fig. *htt-1* functions independently of the DAF-16, DKF-2, EGL-30, FSHR-1, and PMK-1 pathways to promote survival during PA14 infection.** (A) Percent survival of wild-type (WT) and *htt-1(tm8121)* mutant animals with empty vector L4440 or *daf-16* RNAi during PA14 infection (25-35 animals per replicate, $n = 3$). (B) Percent survival of WT and *htt-1(tm8121)* mutant animals with empty vector L4440 or *dkf-2* RNAi during PA14 infection (25-35 animals per replicate, $n = 3$). (C) Percent survival of WT and *htt-1(tm8121)* mutant animals with empty vector L4440 or *egl-30* RNAi during PA14 infection (25-35 animals per replicate, $n = 4$). (D) Percent survival of WT and *htt-1(tm8121)* mutant animals with empty vector L4440 or *fshr-1* RNAi during PA14 infection (25-35 animals per replicate, $n = 2$). (E) Percent survival of WT and *htt-1(tm8121)* mutant animals with empty vector L4440 or *pmk-1* RNAi during PA14 infection (25-35 animals per replicate, $n = 3$). ns (not significant) $p > 0.05$, **** $p \leq 0.0001$ as determined by Log rank test. Detailed statistical analyses are provided in the S1 Table.

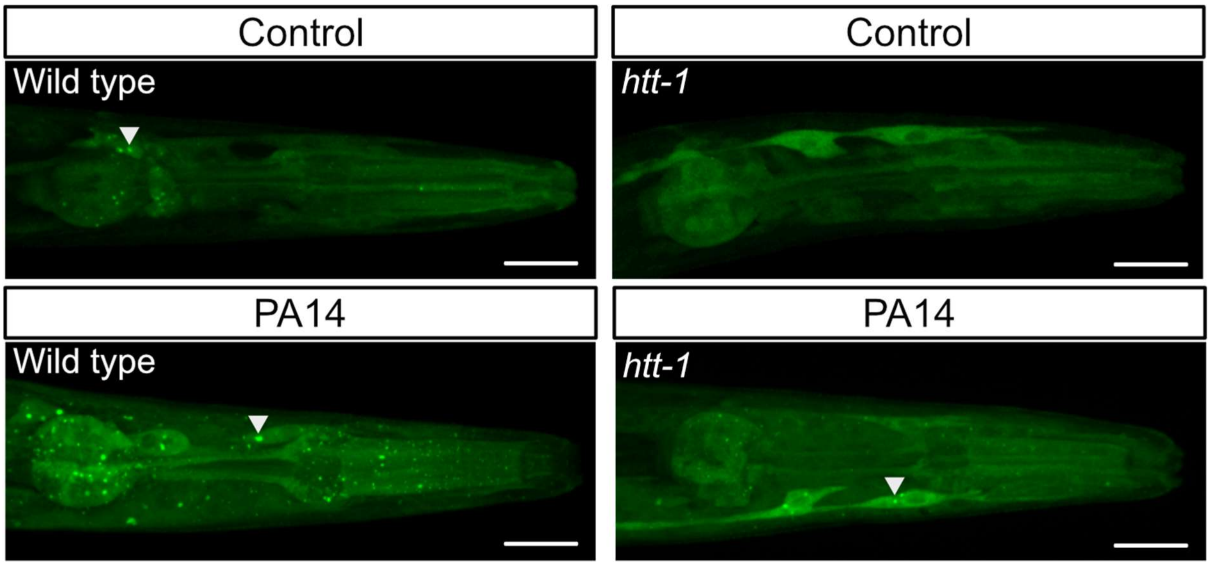


**S5 Fig. *htt-1* mediates PA14 infection-induced autophagy in the head region of *C. elegans.*** Confocal images of GFP::LGG-1 in wild-type N2 and *htt-1(tm8121)* mutant animals. Arrowheads indicate GFP::LGG-1 foci. Scale bars: 20 μm.

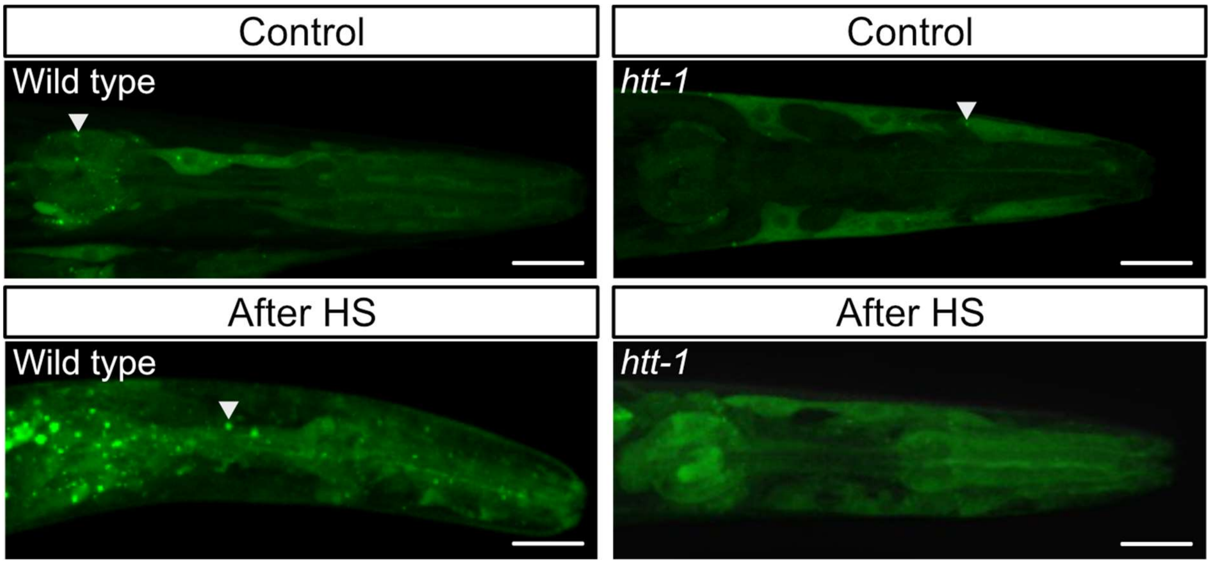


**S6 Fig. *htt-1* mediates heat shock-induced autophagy in the head region of *C. elegans.*** Confocal images of GFP::LGG-1 in wild-type N2 and *htt-1(tm8121)* mutant animals. Arrowheads indicate GFP::LGG-1 foci. Scale bars: 20 μm.

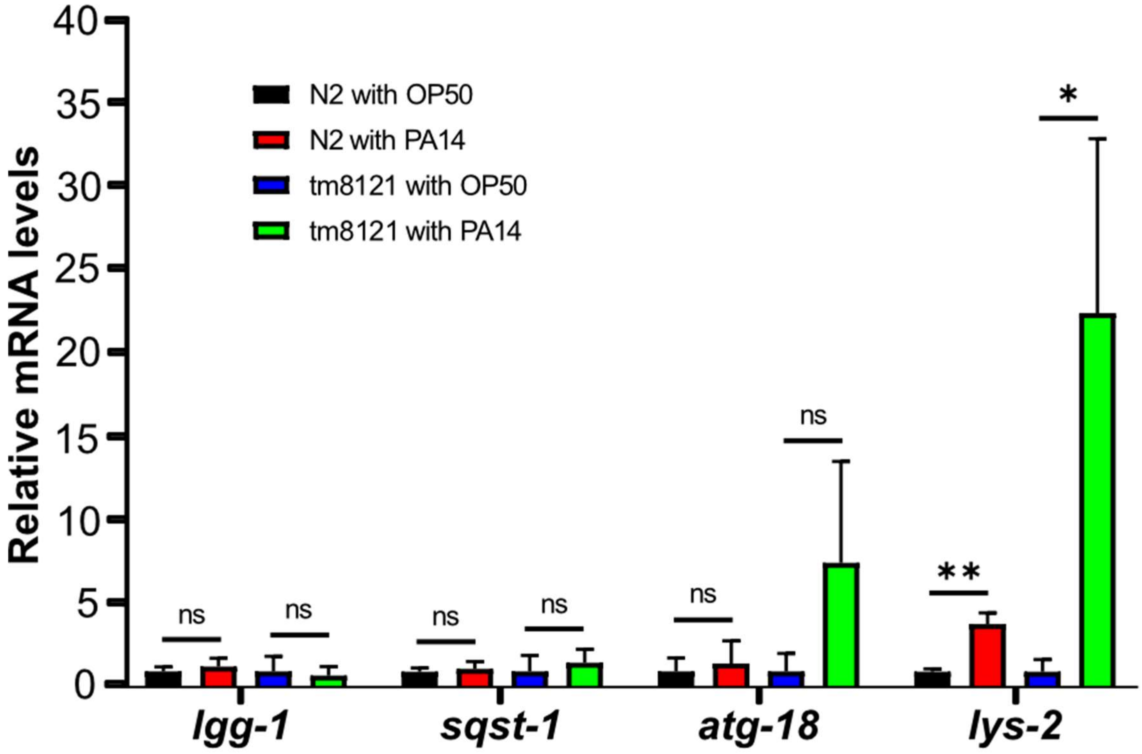


**S7 Fig. Relative mRNA levels of autophagy and lysozyme genes in wild-type N2 and *htt-1(tm8121)* mutant animals cultured on OP50 or PA14 at 25°C for 12 hours.** The mRNA levels of *lgg-1, sqst-1, atg-18,* and *lys-2* were measured by RT-qPCR with *act-1/3* mRNA as an internal control. The fold changes relative to wild-type N2 animals fed by OP50 were shown ($n = 3$). Error bar represent SD. ns (not significant) $p > 0.05$, * $p \leq 0.05$, ** $p \leq 0.01$ as determined by unpaired t test. Detailed statistical analyses are provided in the S5 Table.

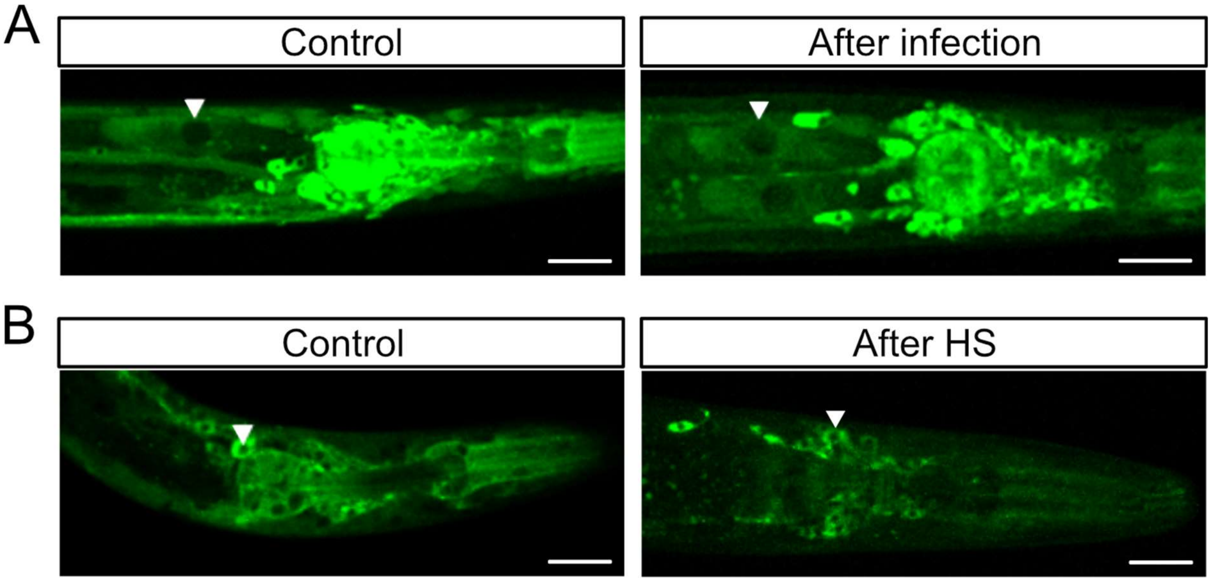


**S8 Fig. HTT-1 remains cytoplasmic.** Confocal images of HTT-1::GFP in wild-type N2 (A) with OP50 and PA14 infection for 4 hours, and *C. elegans* (B) after a 1-hour heat shock at 35°C followed by 1-hour recovery at 20°C. Arrowheads indicate nuclei. Scale bars: 20 μm.
